# DiMSum: an error model and pipeline for analyzing deep mutational scanning data and diagnosing common experimental pathologies

**DOI:** 10.1101/2020.06.25.171421

**Authors:** Andre J. Faure, Jörn M. Schmiedel, Pablo Baeza-Centurion, Ben Lehner

## Abstract

Deep mutational scanning (DMS) enables multiplexed measurement of the effects of thousands of variants of proteins, RNAs and regulatory elements. Here, we present a customizable pipeline – DiMSum – that represents an end-to-end solution for obtaining variant fitness and error estimates from raw sequencing data. A key innovation of DiMSum is the use of an interpretable error model that captures the main sources of variability arising in DMS workflows, outperforming previous methods. DiMSum is available as an R/Bioconda package and provides summary reports to help researchers diagnose common DMS pathologies and take remedial steps in their analyses.

## Background

Deep mutational scanning (DMS), also known as massively parallel reporter assays (MPRAs) and Multiplex Assays of Variant Effect (MAVEs), enables parallel measurement of the effects of thousands of mutations in the same experiment[1, 2]. In a basic DMS experiment a library of sequence variants is constructed and deep sequencing before and after selection for an *in vitro* or *in vivo* activity is used to quantify the relative activity (‘molecular fitness’) of each genotype. Beyond assaying point mutations, the high-throughput nature of DMS facilitates the comprehensive study of combinations of mutations and their genetic interactions (epistasis) where fitness effects of individual mutations depend on the presence of other (background) mutations[3]. The resulting fitness landscapes are informative of protein[4–6], RNA[7–9] and regulatory element[10–18] function, and have provided mechanistic insight into biological processes including the regulation of gene expression[10,19], protein-protein interactions[20], alternative splicing[21,22] and molecular evolution[7]. Deep mutational scans have the potential to improve human variant annotation[23,24] and protein and RNA structure determination[25–27]. In recognition of the growing number and importance of DMS assays in biomedical research, a dedicated platform for sharing, accessing and visualizing these datasets has recently been launched[28].

A key feature of a DMS experiment is that it preserves the link between quantitative phenotypic effects and their underlying causal genotypes measured for many variants simultaneously (Figure 1a). The three main steps can be summarized as follows: (1) construction of a library of DNA variants corresponding to the assayed biomolecule (genotype), (2) selection (or separation) of variants according to a given molecular function (phenotype), and (3) quantification of the variant abundances before and after selection by DNA sequencing (measurement), which is either done by counting sequencing reads of variants directly or unique barcodes previously linked to them[29–33]. A fitness score for each variant is then calculated by comparing its relative abundance (with respect to a reference sequence e.g. wild-type) before and after selection. Moreover, often multiple independent biological replicates of the experiment are performed to help estimate the error of variant fitness scores, that is, a measure of fitness score reliability.

**Figure 1.**
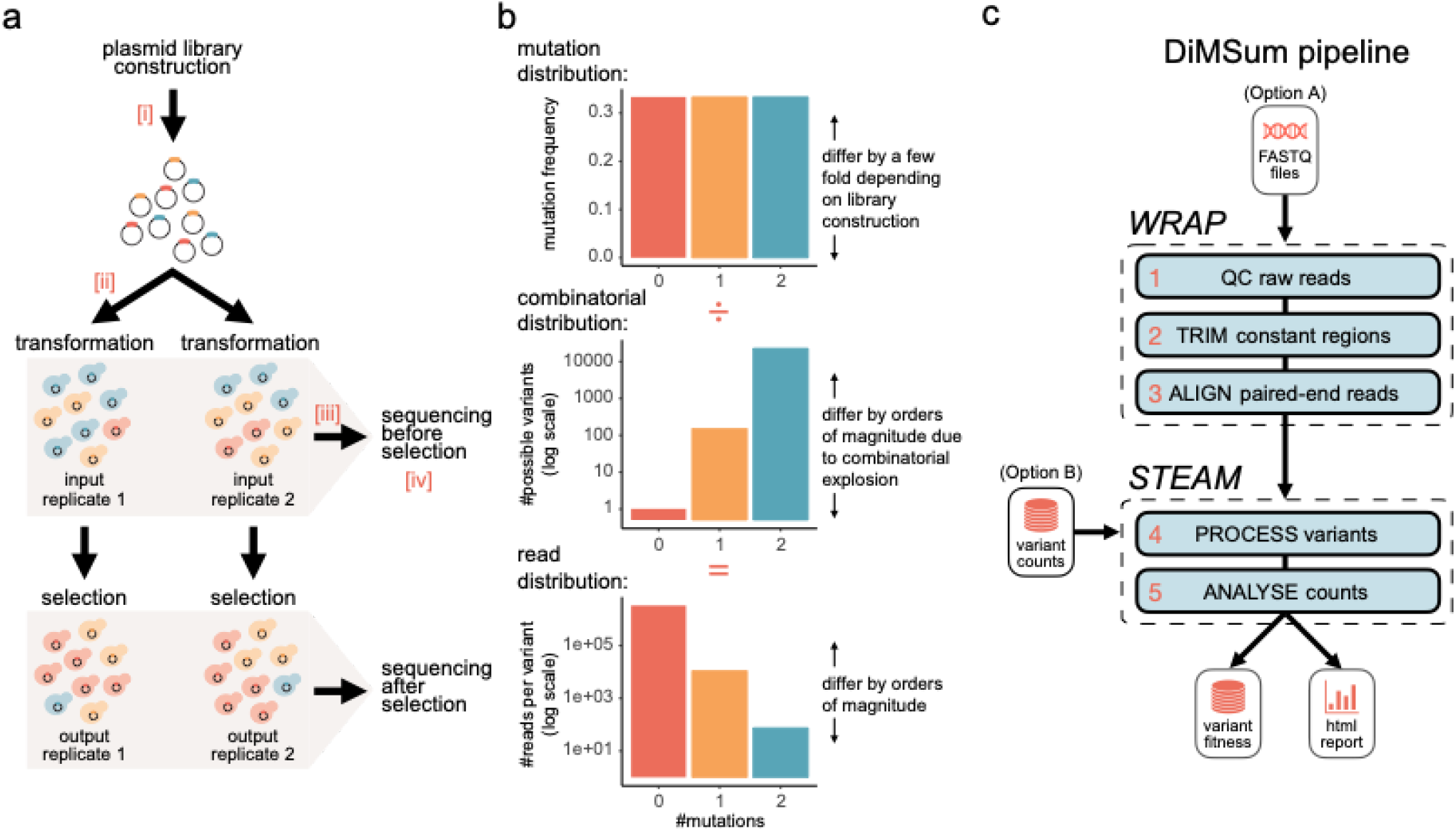
Schematic overview of a minimal DMS experiment and the DiMSum pipeline. **a.** Schematic of a basic, plasmid-based microbial growth DMS experiment: (1) construction of a plasmid-library of mutant variants and independent transformation or integration of plasmid-library into host cells, (2) exposure of cell population to selective conditions, (3) high-throughput sequencing of samples to obtain variant counts before and after selection, which are used to derive fitness estimates for each variant. Indicated are steps at which bottlenecks could arise, potentially restricting variant pool size or complexity (red roman numerals): [i] inefficient library construction (“Library bottleneck”), [ii] inefficient plasmid transformations (“Replicate bottleneck”) and [iii] inefficient DNA extraction (“DNA extraction bottleneck”). Unforeseen bottlenecks can lead to over-sequencing [iv] of variant pools and thus under-estimation of the errors associated with fitness scores or even appearance of sequencing counts for variants not contained in the original variant pool. **b.** DMS experiments typically have a hierarchical abundance structure, where variants with more mutations are orders of magnitude less abundant than the wild-type sequence or single mutants. **c.** DiMSum flow chart. The *WRAP* module performs low-level processing of raw DNA sequencing reads to produce sample-wise variant counts. The *STEAM* module transforms the resulting counts to estimates of variant fitness and associated error. See Supplementary Figures 1-6 for example report plots.

Several software packages have been developed to simplify and standardize the calculation of fitness scores for each variant from deep sequencing data[34,35], including the estimation of errors for these fitness scores[36]. Unbiased estimation of fitness score reliability is crucial for the interpretation of DMS experiments, for example when assessing the effects of a variant in a disease gene, and more generally for all kinds of hypothesis testing and when assessing genetic interactions.

The large-scale construction and high-throughput readout of thousands to hundreds of thousands of variants at once can, however, complicate basic quality control and identification of potential error sources and artefacts arising in DMS workflows. On the one hand, the many experimental steps of a DMS workflow can contribute errors to the final fitness measurements, especially when ‘bottlenecks’ restrict the variant pool at certain steps in the workflow (Figure 1a). On the other hand, libraries with a hierarchical variant abundance structure, arising from the combinatorial explosion of variants with multiple mutations (Figure 1b), leads to distinct sources of error differentially affecting specific subsets of variants (see below). Moreover, the hierarchical variant abundance structure in combination with the typically low complexity of the genotype pool can lead to artefacts introduced by sequencing errors[37,38].

To tackle these issues, we developed DiMSum, a pipeline that allows the end-to-end processing of DMS datasets using an interpretable model for the magnitude and sources of errors in fitness score. The workflow is freely available as an R/Bioconda package (DiMSum) that represents a complete solution for obtaining reliable variant fitness scores and error estimates from raw sequencing files.

## Results and Discussion

### Overview of DiMSum pipeline

The DiMSum pipeline is implemented as an R/Bioconda package and command-line tool that can be easily configured to handle a variety of DMS experimental designs (see Methods). The pipeline is organized in two separate modules (Figure 1c): *WRAP* processes raw read (FASTQ) files to produce sample-wise variant counts, and *STEAM* uses these sample-wise variant counts to estimate variant fitness scores and their measurement errors.

DiMSum *WRAP* performs the following sequence processing steps: (1) assessment of raw read quality using FastQC[39], (2) error-tolerant removal of constant regions (not subjected to mutagenesis but required for primer binding and isolation/amplification of variables regions) using cutadapt[40] and (3) alignment and filtering of paired-end reads in a base quality-aware manner using VSEARCH[41] if required. DiMSum *STEAM* accepts a table of counts, (4) isolates substitution variants of interest and then (5) performs statistical analyses to obtain associated fitness scores and error estimates. Briefly, an error model is fit to a high confidence subset of variants to determine count-based, additive and multiplicative errors of variant fitness scores for all replicates (see below).

To increase flexibility, *WRAP* or *STEAM* can each be run in stand-alone mode if desired, e.g. to obtain fitness scores from a user-generated table of variant counts (Figure 1c, ‘Option B’) or to obtain sample-wise variant counts for a custom downstream analysis (Figure 1c, ‘Option A’). A detailed R markdown report – viewable with any web browser – including summary statistics, diagnostic plots and analysis tips is also generated.

### Estimates of variant fitness scores and associated errors

DiMSum calculates variant fitness scores as the natural logarithm of the ratio between sequencing counts in a replicate’s output and input samples relative to the wild-type variant. It then uses replicate-specific error estimates to produce a weighted average of fitness scores across replicates for each variant.

DMS experiments are typically replicated to judge the reliability of fitness score estimates due to random variability in the workflow. However, the number of replicates performed is usually low (e.g. 3 to 6), and estimates of measurement errors on a variant-by-variant basis can thus lack statistical power. DiMSum instead estimates measurement errors of fitness scores by sharing information across all assayed variants to increase statistical power (Figure 2, see Methods for full detail).

**Figure 2.**
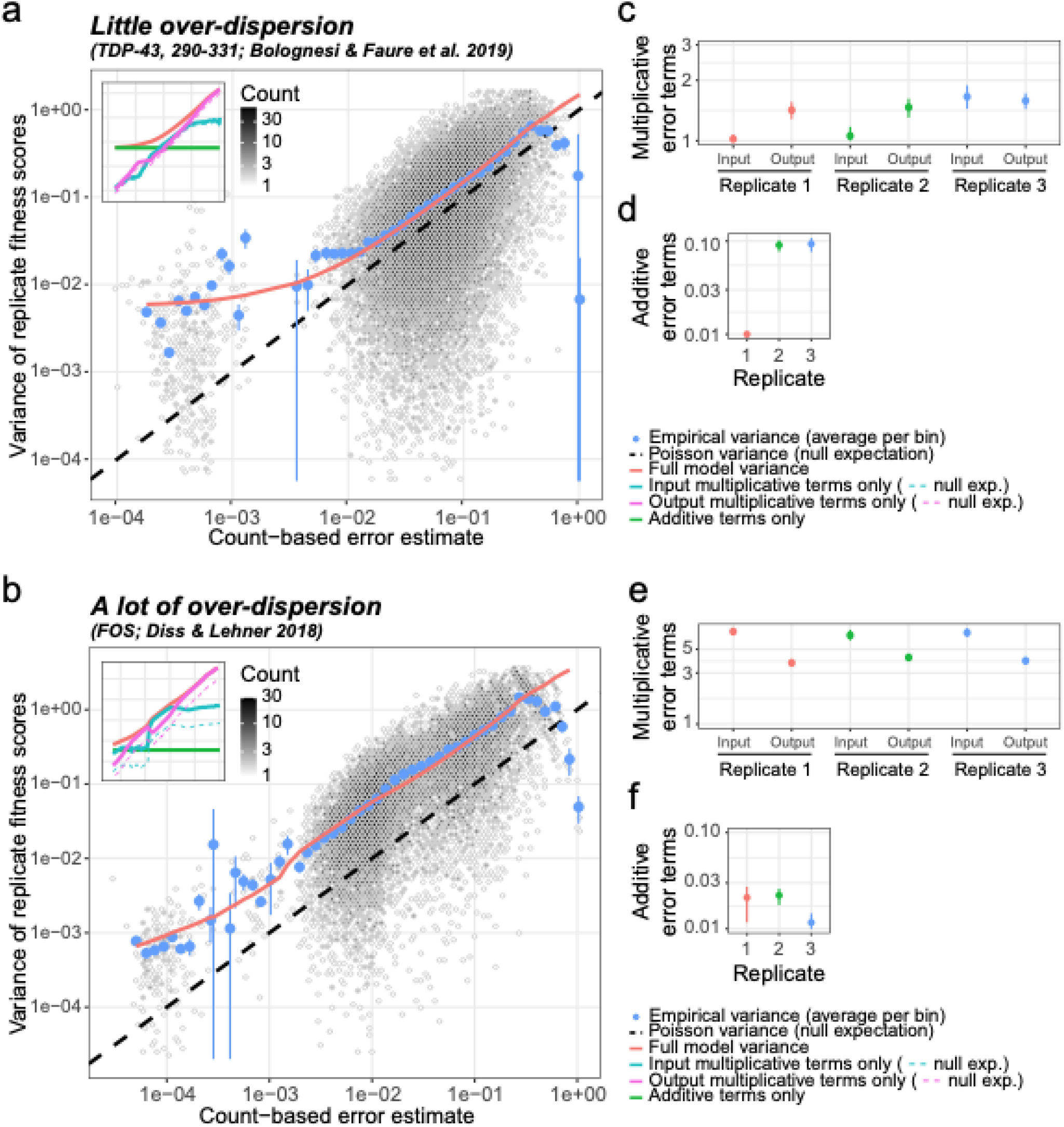
DiMSum error model estimates multiplicative and additive error sources in fitness scores. **a**. Empirical variance of replicate fitness scores as a function of error estimates based on sequencing counts under Poisson assumptions in a deep mutational scan of TDP-43 (positions 290-331)[6]. Empirical variance (blue dots show average variance in equally-spaced bins, error bars indicate *avg. variance* × (1 ± 2/*#variants per bin*)) is over-dispersed compared to baseline expectation of variance being described by a Poisson distribution (black dashed line). The bimodality of the count-based error distribution results from the relatively low number of single nucleotide mutants which have high counts (thus low count-based error) and the many double nucleotide mutants which have low counts (thus higher count-based error). The DiMSum error model (red line) accurately captures deviations of the empirical variance from Poisson expectation. Inset: bold cyan and magenta lines indicate multiplicative error term contributions to variance corresponding to Input and Output samples respectively (dashed thin lines give Input or Output sample contributions to variance if multiplicative error terms were 1). The horizontal green line indicates the additive error term contribution. The red line indicates the full DiMSum error model. **b**. Same as panel **a** but for a deep mutational scan of FOS[20] that shows more over-dispersion. **c-f**. Multiplicative (panels **c** and **e**) and additive (in s.d. units, panel **d** and **f**) error terms estimated by the error model on the two datasets. Dots give mean parameters, error bars 90% confidence intervals.

We assume that the error in fitness scores is, to a first approximation, primarily arising due to the finite sequencing counts and thus variants with similar counts in input and output samples should have similar measurement errors[42,43]. If error was purely arising due to sampling of variant frequencies by sequencing, the error could be well approximated by a Poisson distribution, with variance equal to the mean[44]. However, count data have been found to often be over-dispersed compared to this base-line Poisson expectation[45,46]. To account for such over-dispersion, we introduce additive and multiplicative modifier terms of the base-line error, which has been shown to accurately describe variability in transcriptomic count data[47–49].

Multiplicative error terms modify the overall error proportional to the error resulting from a variant’s sequencing counts, and likely describe error sources in workflow steps linked to sequencing (see below for a discussion of potential sources and experimental remedies). Across different DMS datasets, we find such multiplicative error terms to range from one all the way to more than 100, suggesting that over-dispersion can be a grave issue in DMS experiments (Figure 2, Table 2).

Additive error terms are independent of a variant’s sequencing read counts, thus affecting all variants to the same extent, which we attribute to variability arising from differential handling of replicate selection experiments (see below). Additive error terms are typically small (*s.d*. < 10%) and therefore only become apparent in variants that have small errors from sequencing counts (those with many counts), constituting a lower error limit (Figure 2, Table 2).

We assume that both multiplicative and additive error terms can differ between replicates but are the same for all variants in each replicate; our error model therefore has 3*n* parameters (where n is the number of replicates), which are estimated by minimizing the squared difference between the empirical and model-predicted variance of fitness scores across replicates for all variants simultaneously (see Methods).

Manipulating a DMS dataset to artificially increase either multiplicative error terms or additive error terms in one replicate, suggests that the DiMSum error model is capable of accurately estimating the magnitude of the error model terms (Supplementary Figure 7).

### Error model benchmarking

To benchmark the error model, we performed leave-one-out cross validation on published DMS datasets. Here, error model parameters were trained on all but one experimental replicate of a dataset. The resulting error estimates were used to judge whether the fitness scores of variants differ between the training replicates and the held-out test replicate. We find fitness scores differences between training and test replicates are normally distributed with the magnitude predicted by the error model (Figure 3a). Consequently, when testing for significant differences between the training and test replicates (using a z-test), P-values are uniformly distributed (Figure 3b), as would be expected for replicates from the same experiment and indicating that the model correctly controls the type I error rate (rate of false positives).

**Figure 3.**
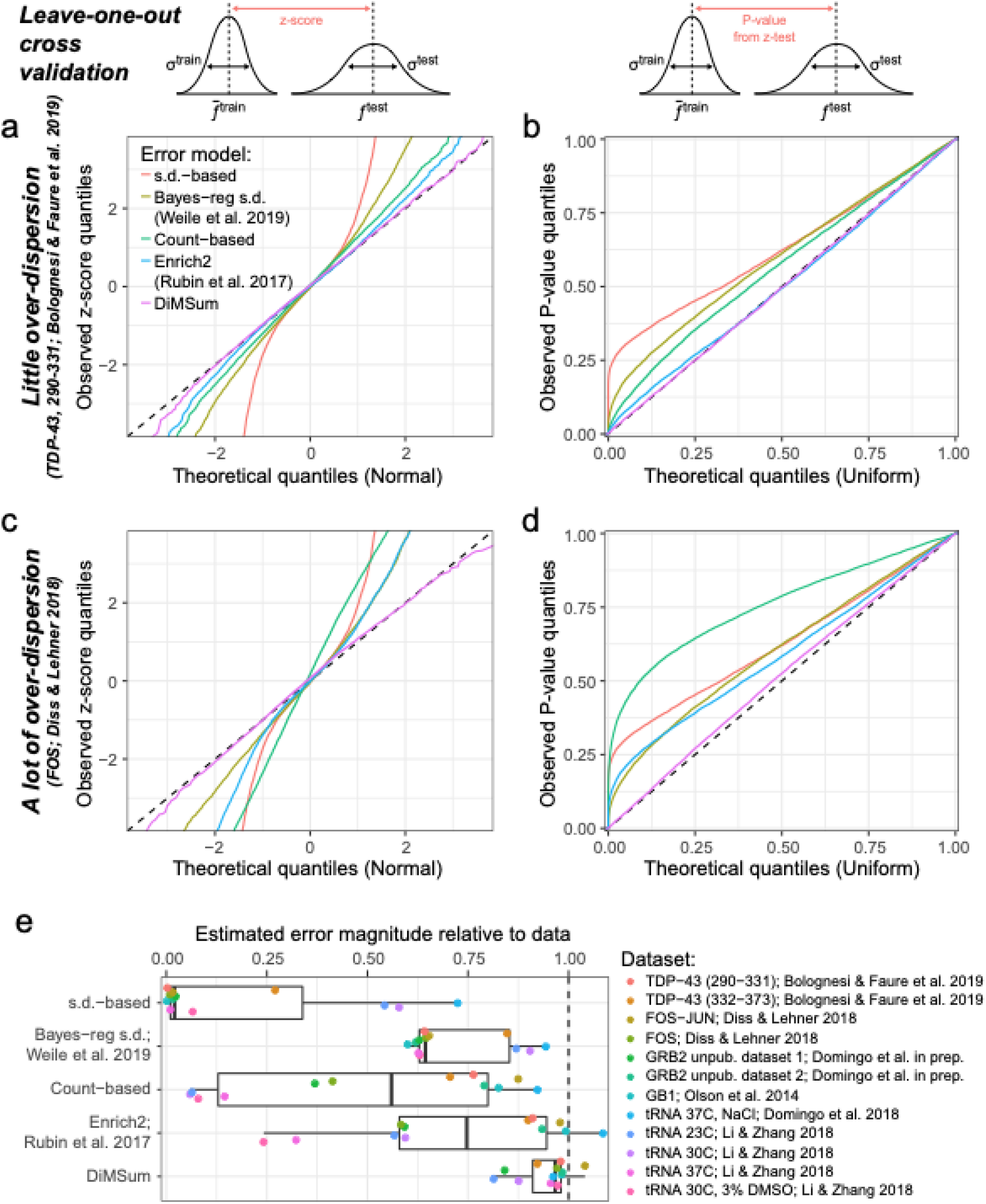
DiMSum error model performance. Leave-one-out cross validation to test error model performance. In turn, error models are trained on all but one replicate of a dataset and z-scores of the differences in fitness scores between the training set 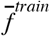 and the remaining test replicate *f^test^* are calculated (i.e. fitness score differences normalized by estimated error in training set σ^*train*^ and test replicate σ^*test*^. Importantly, σ^*test*^ is estimated from error model parameters fit only on the training set replicates). Because fitness scores from replicate experiments should only differ by random chance, if the error models estimate the error magnitude correctly, z-scores should be normally distributed, and corresponding p-values from a z-test should be uniformly distributed. The tested error models are described in Results and Methods sections. **a & c.** Quantile-quantile plots of z-scores in TDP-43 290-331 library (**a**) and FOS library (**c**) compared to expected Normal distribution. **b & d**. Quantile-quantile plots of P values from two-sided z-test in TDP-43 290-331 library (**a**) and FOS library (**c**) compared to expected uniform distribution. **e**. Estimated error magnitude relative to the differences observed between replicate fitness scores in twelve DMS datasets in leave-one-out cross validation (see Methods). Relative error magnitude = 1 means the estimated magnitude of errors fits the data, Relative error magnitude < 1 means the estimated errors are too small. Boxplots indicate median, 1st and 3rd quartile (box) and whiskers extend to 1.5 × interquartile range.

We find that the DiMSum error model accurately estimates errors in fitness scores across twelve published DMS datasets that display various degrees of over-dispersion (Figure 3, Table 2). Moreover, errors are accurately estimated no matter whether they are driven by low sequencing counts (variant with low counts, often higher-order mutants) or whether they appear to be independent of sequencing counts (variants with high counts, such as single mutants), suggesting that both multiplicative and additive error terms help to accurately model error sources in DMS experiments (Supplementary Figure 8).

We compared the DiMSum error model performance to several popular alternative approaches that have previously been used to model error in DMS data (see Table 1). We note that this is not an exhaustive comparison against all statistical models previously used before to estimate measurement errors in DMS datasets. The chosen alternative approaches differ in whether they estimate errors for each variant from the observed variability of fitness scores or the sequencing counts, or a combination thereof, and how much information sharing across variants they allow.

**Table 1:** Benchmarked statistical models to estimate measurement error in fitness scores

| Error model type | Based on | Additional input? | Information sharing across variants? | Model parameters |
| --- | --- | --- | --- | --- |
| variance-based | Empirical variance of replicate fitness | **No** | **No** | 0 |
| Bayesian regularization of variance [51] | Empirical variance of replicate fitness | **Yes:** Bayesian prior from regression on seq. counts | **Yes:** Prior estimated across all variants | 3 |
| count-based | Sequencing counts follow Poisson distribution | **No** | **No** | 0 |
| Enrich2 [36] | Sequencing counts follow Poisson distribution | **Yes:** Mixed-effects from empirical variance | **No** | # variants |
| DiMSum | Sequencing counts follow Poisson distribution | **Yes:** Modifier terms from empirical variance | **Yes:** Replicate-specific modifier terms estimated across all variants | 3 x # replicates |

On the one hand, several studies have used the empirical variance of fitness scores across replicates to calculate errors for each variant individually[8,9,20,22,31,50]. Such error estimates are under-powered due to the typically low number of replicates in DMS studies, resulting in errors that are too large for some variants but too small for others. The latter results in an inflation of type I errors (Figure 3b,d, ‘s.d.-based’). Error estimates improve with increasing number of replicates, but type I error inflation persists even for a DMS dataset with 6 replicates (Figure 3e, Domingo et al., 2018[7]).

Building on this empirical variance approach, Weile et al.[51] used a Bayesian regularization of the empirical variance proposed by Baldi and Long[52], which uses a linear regression estimate of empirical variance across all variants as a prior. We find that this approach improves over only using the empirical variance to calculate errors, but still leads to inflation of type I errors (Figure 3, ‘Bayes-reg s.d.’).

On the other hand, several studies have assumed that errors can be modeled by a Poisson process based on a variant’s sequencing counts[5–7,53]. Not unexpectedly, the performance of the ‘Poisson-based’ approach depends on the over-dispersion of the data. It works well for datasets with little systematic over-dispersion, but fails dramatically in those cases where the DiMSum error model estimates high multiplicative or additive errors (Figure 3 and Supplementary Figure 8, ‘Count-based’).

Enrich2[36] uses a random-effect model to account for overdispersion over and above the count-based Poisson expectation on a variant-by-variant basis. In short, variant-specific random effect terms increase the modelled error towards the empirical variance if it is larger than the count-based Poisson expectation. While this leads to accurate error estimates in datasets with little systematic over-dispersion (small multiplicative error terms, Figure 3a, c, ‘Enrich2’), the underpowered estimation of the variant-specific random-effect terms leads to an inflation of type I errors in those datasets with systematic over-dispersion, similar to the other approaches based on variant-specific empirical variances (Figure 3b, d, e, ‘Enrich2’).

In summary, the DiMSum error model captures the major error sources arising in DMS workflows and improves in accuracy over previous approaches, while needing fewer replicate experiments and having fewer, but interpretable model parameters.

DiMSum provides diagnostic plots similar to Figure 3a,c to help judge whether errors have been accurately modeled. Failure of the model to accurately estimate the errors suggests shortcomings, potentially due to systematic error sources in the DMS workflow which cannot be accounted for by the error model, urging further action by the user (see below).

### Potential sources of increased error in fitness score estimates

In what follows, we provide suggestions for error sources that might be captured by DiMSum’s additive and multiplicative error model terms as well as error sources that cannot be captured by the error model and how their impact on DMS experiments can be minimized.

**Additive error terms** are independent of variant read counts and therefore likely result from differential handling of replicate selection experiments. Because these error terms are typically small compared to errors resulting from low sequencing counts, they most often only affect fitness score estimates of very abundant variants, such as single mutants of the wild-type sequence in question. However, if such highly abundant variants are of interest, increasing sequencing coverage will not lead to reductions in the measurement errors of their fitness scores. DiMSum performs a simple scale and shift procedure to minimize inter-replicate differences in fitness score distributions prior to estimating error model parameters, therefore minimizing additive error terms that arise from linear differences between replicate selection experiments (see Methods and Supplementary Figure 4b,c). Additional mitigation strategies to reduce additive error contributions should focus on streamlining the handling of replicate samples through the workflow (e.g. using master mixes, increasing pipetting volumes, reducing time-lags in time sensitive steps) as well as increasing the number of replicate experiments[53], even at similar overall sequencing coverage, as this will lead to a reduction of errors for variants that are dominated by sequencing-independent errors due to the weighted averaging of fitness scores across replicates.

**Multiplicative error terms** increase variants’ errors by a multiple of their sequencing count based error estimate. Potential error sources are thus likely linked to the sequencing steps in the DMS workflow, in particular related to the start of the selection step, DNA extraction from input and output samples as well as the subsequent PCR amplification for sequencing library construction.

First, consider a bottleneck at the DNA extraction step, which arises if the number of unique DNA molecules extracted from the input/output samples does not exceed the number of molecules that are subsequently sequenced, i.e. the extracted variant pool is ‘over-sequenced’. This restriction in the numbers of variant molecules along the workflow will introduce additional random variability in variant frequencies that significantly contribute to – or even dominates – the overall count-based error, and errors calculated solely from the number of downstream sequenced molecules will thus be an underestimate of the true error.

In addition, Kowalsky et al.[54] found that PCR amplification protocols for sequencing library construction can introduce additional random variability to variant frequencies. Using our DiMSum error model, we find that multiplicative errors differ five-fold between the three PCR protocols tested (see Methods), thus showing that multiplicative errors can arise during the PCR amplification steps of the DMS workflow.

Lastly, another source of multiplicative errors that can potentially arise in input samples is a bottleneck at the start of the selection experiment. Here, if the number of variant molecules used to start the selection is similar to or smaller than the number of variant molecules extracted and sequenced from the input sample, this will randomly alter true variant frequencies at the start of the selection with error magnitudes on the order of or even larger than the error due to sequencing a finite subset of variant molecules.

For example, we recently performed a deep mutational scan of part of the protein GRB2 (Domingo et al., manuscript in preparation), for which the error model indicated a six-fold multiplicative error in the input replicates. A similar error was not observed in a second, related deep mutational scan for the same protein, suggesting a technical bottleneck specific to the input library preparation in the first experiment.

To minimize multiplicative error sources, thus reduce measurement errors and ultimately save sequencing costs, DMS workflows should ensure an excess of variant molecules (~5x-10x) are used in all experimental steps upstream of the sequencing step[55] and PCR amplification protocols are optimized[54]. Additionally, sources of multiplicative errors due to bottlenecks at the DNA extraction step and other downstream steps, but not during the selection experiment, should be detectable (and correctable) if using unique molecular identifiers (UMIs) ligated to variant molecules during PCR-based sequencing library preparation[31,56,57].

#### Systematic error sources

Apart from sources of increased measurement error due to random error in DMS workflows, there are potentially also sources of systematic error that the DiMSum error model cannot account for and which might therefore inflate error or bias fitness scores in undetectable ways.

One potential source of systematic errors is (non-linear) differences in the replicate selection experiments. For example, we recently used DMS to quantify the toxicity of variants of deTDP-43 when expressed in yeast in which we mutagenized two sections of the C-terminal prion-like domain[6]. Variants displayed a range of fitness values relative to the wild-type sequence, both detrimental and beneficial. Importantly, one replicate experiment showed a marginal fitness distribution whose shape differed from those of three other replicate experiments. In particular, non-toxic mutant variants were limited in how much faster they could grow compared to wild-type TDP-43, which perhaps resulted from nutrient limitation during the selection experiment (Supplementary Figure 4c). Such non-linear effects that only affect a subset of variants (e.g. beneficial variants) cannot be corrected for with simple linear normalization schemes (e.g. DiMSum’s shift and scale normalization procedure) and will introduce systematic errors that the error model cannot adequately describe, thus potentially leading to biased fitness estimates as well as incorrect estimates of errors (Supplementary Figure 6e,f). Thus, systematic differences in replicate selection experiments identify the need for better normalization strategies or exclusion of affected replicates, as we decided for the TDP-43 replicate[6].

In summary, the DiMSum error model and diagnostic plots can also serve to judge and improve the experimental workflow and downstream analyses of DMS experiments.

### Diagnosing sources of systematic errors in DMS workflows

The particular combination of low genotype complexity and hierarchical abundance structure in DMS experiments (Figure 1b) can lead to issues arising from sequencing errors. On the one hand, sequencing errors in reads of highly abundant variants can contribute counts to closely related, but low abundant, variants[37,38]. That is, sequencing errors in wild-type reads will contribute counts to single mutant variants, and sequencing errors in single mutant variants will contribute counts to double mutant variants and so on. DiMSum displays estimates of this sequencing error-induced ‘variant flow’ in diagnostic plots of marginal count distributions to give the user an estimate of what fraction of reads of a set of mutants might be caused by sequencing errors (Figure 4a, left column). Mitigation strategies to lower the fraction of reads per variant from sequencing errors include using higher minimum base quality (Phred score) thresholds, using paired-end sequencing to decrease the number of base call errors, or circumventing these issues altogether by using highly complex barcode libraries that are linked to variants[38].

**Figure 4.**
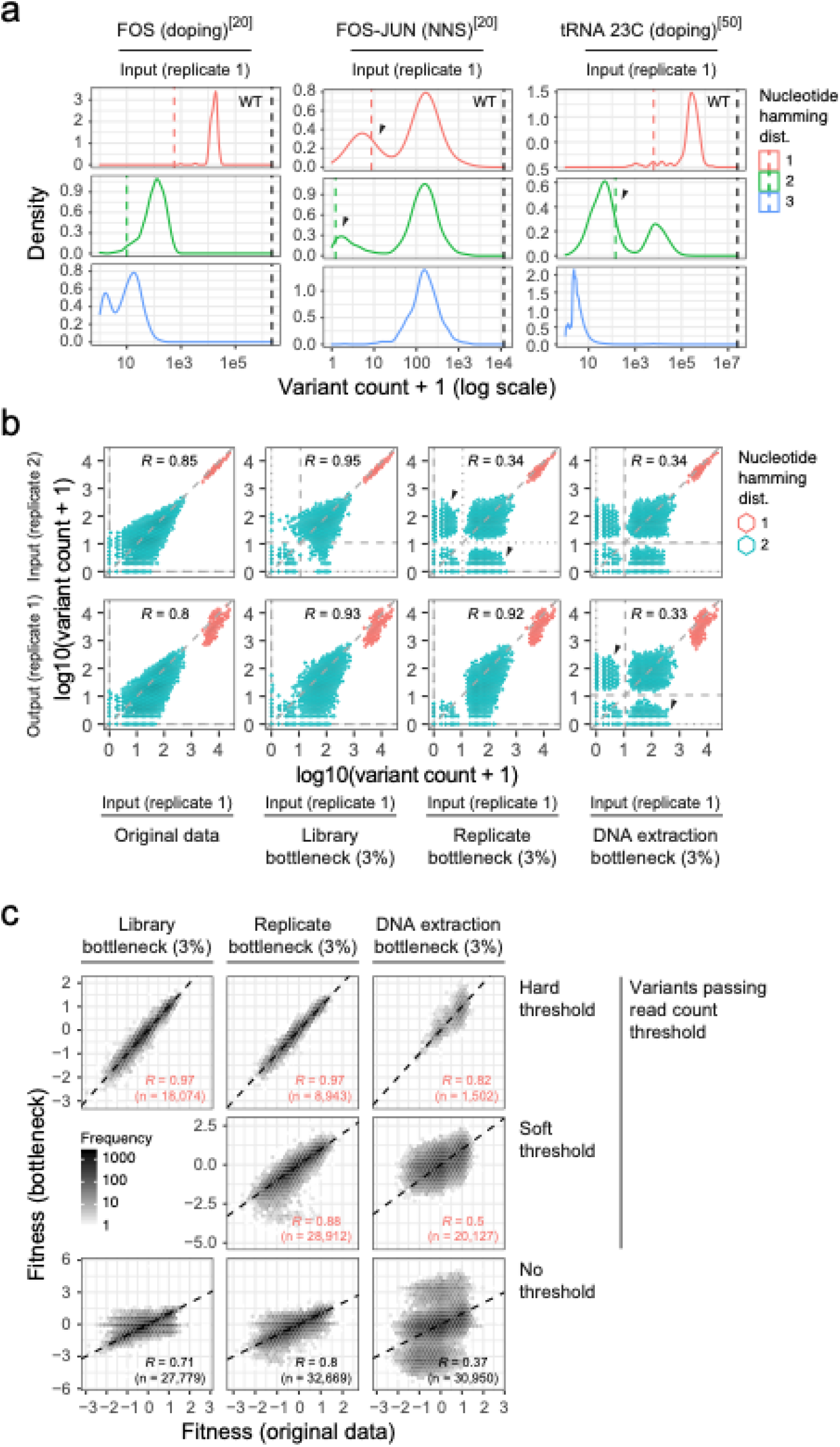
Effects of bottlenecks on variant count distributions and fitness scores. **a**. Input sample count distributions of previously published DMS experiments[20,50]. For FOS and FOS-JUN datasets, counts of single AA variants with one, two or three nucleotide substitutions in the same codon are shown. For the tRNA dataset, all variants with one, two or three nucleotide substitutions are shown. Wild-type counts are indicated by the black dashed line. Expected count frequencies purely due to sequencing errors are indicated by red and green dashed lines for single and double nucleotide substitution variants, respectively. Black arrow indicates sets of variants that have likely not been assayed but whose sequencing reads are arising due to sequencing errors. **b**. Simulation of bottlenecks at various steps of the DMS workflow based on a previously published DMS dataset[6]. Scatterplots show Input and Output sample counts for variants with one or two nucleotide substitutions in the original data or after simulating 3% library, replicate or DNA extraction bottlenecks (from left to right). Hexagon color indicates number of nucleotide substitutions and fill number of variants per 2d bin (see legend). Black arrows indicate sets of double nucleotide variants whose sequencing reads solely originate from sequencing errors. Dotted (or dashed) horizontal/vertical lines indicate soft (or hard) variant count thresholds used in downstream DiMSum analyses (see panel **c**). **c**. Comparison of fitness scores from simulated datasets with (y-axis) or without (x-axis) the indicated bottlenecks. Variants are categorized by their robustness to filtering with hard (variants have to appear above threshold in all replicates) or soft thresholds (variants have to appear above threshold in at least one replicate) of 10 read counts. For the DNA extraction bottleneck, read count thresholds were also applied to output samples. Pearson correlation coefficients are indicated. The dashed line indicates the relationship y=x. Note that correlation coefficients are lower for soft than hard thresholds, because a subset of variants has fewer replicate measurements.

On the other hand, a potential pitfall linked to the combination of low genotype complexity, hierarchical abundance structure and sequencing errors in DMS experiments is to mistake sequencing reads purely arising from sequencing errors for the presence of a variant in the assayed genotype pool. That is, at deep enough sequencing coverage, reads for any low-order nucleotide mutant variant will appear in the sequencing record, even if the variant was not actually present in the experiment.

Consider two examples from published DMS experiments. The first example of a DMS experiment in which NNS (N=A,T,C or G, S=C or G) saturation mutagenesis was used to introduce individually mutated codons into the wild-type sequences of FOS and JUN[20]. Variants that have one mutated codon show a bimodal count distribution in the input samples (Figure 4a, middle column). Variants in the higher peak have similar read counts no matter whether one, two or three nucleotides were mutated, consistent with NNS mutagenesis operating on the codon level and the number of mismatch base-pairs having little impact on mutation efficacy. In contrast, read counts for variants in the lower peaks show a dependency on the number of nucleotides mutated and coincide with DiMSum’s estimate for sequencing error-induced variant flow. The second example is from a DMS experiment in which doped oligonucleotide synthesis was used to introduce nucleotide mutations into a tRNA[50]. Variants with one or two mutated nucleotides show a bimodal count distribution in the input samples (Figure 4a, right column). The read counts for variants in the upper peaks depend on the number of nucleotides mutated, consistent with read counts per variant being strongly affected by the combinatorics of mutational space (Figure 1b). The read counts for variants in the lower peaks also depend on the number of nucleotides mutated and coincide with DiMSum’s estimate for sequencing error-induced variant flow.

Are variants in the lower read count peaks of these experiments really not present in the variant libraries before sequencing? And at which steps in the DMS workflow were the variants lost? Potential bottlenecks (or inefficacies) might arise during library construction, transfer of the library into the assay cell population or at subsequent DNA extraction and sequencing library preparation steps (Figure 1a).

We find that the comparison of count distributions between sequencing samples can provide additional support to determine whether subsets of variants purely arose from sequencing errors and help to diagnose at which workflow step variants might have been lost, in order to improve future DMS experiments and serve to inform the strategy to avoid systematic errors in fitness calculations for a present dataset. We exemplify this in Figure 4b on simulated bottlenecks in a deep mutational scan of TDP43.

If variants have not been constructed or have been lost at initial library preparation steps and therefore are not present in any replicate experiment, count distributions between replicate input samples should be highly correlated and the same variants should fall into the same peaks of bimodal read count distributions (Figure 4b, ‘library bottleneck’), as is also apparent in the FOS-JUN dataset (Supplementary Figure 9). Variants in the lower peak of the distribution should be discarded from all replicates, e.g. using DiMSums ‘hard’ read count thresholds for variant filtering (Figure 4c), and downstream analysis should proceed as normal, as in the published analysis of the FOS-JUN dataset [20].

In contrast, if variant loss was replicate-specific, e.g. if transformations into replicate cell populations were incomplete, read count distributions should display ‘flaps’ – subsets of variants that appear at high counts in one replicate (variant was assayed) but at low counts in another (variant counts arise solely from sequencing errors) (Figure 4b, ‘replicate bottleneck’). A conservative approach to avoid systematic errors in fitness score calculations is to use ‘hard’ read count thresholds to discard all variants appearing in lower read count peaks in any replicate. Additionally, DiMSum allows the user to choose a ‘soft’ threshold to discard variants only in the replicates where they appear in the low count peaks, therefore allowing their fitness to still be estimated from replicates in which they are actually present, resulting in increased number of variants that can be used for downstream analyses (Figure 4c).

Finally, if variants were lost at the DNA extraction steps, this should not only show up as flaps in count distributions between replicate input samples, but also between input and output samples of the same replicate (Figure 4b, ‘DNA extraction bottleneck’), as it is observed for the tRNA dataset (Supplementary Figure 9). Here, in order to avoid biased fitness estimates, all variants that do not appear in the high read count peak of both input and output samples from the same replicate experiment need to be discarded to avoid systematic errors in downstream analyses. Often fitness differences between variants also result in bimodal output count distribution, meaning that practically it can be hard or impossible to assign whether variants with low counts in output samples are due to low fitness or because they were not assayed. As for the replicate bottlenecks, ‘soft’ thresholds can be used to obtain fitness estimates for all variants that appear in the input and output samples of at least one replicate, therefore increasing the number of variants that can be used for downstream analyses (Figure 4c).

To further illustrate how experimental bottlenecks can adversely affect the conclusions of a study, we evaluated their impact on the central conclusion of a previous publication. We previously showed that the fitness effects of amino acid substitutions in the prion-like domain of TDP-43 are correlated with the increase in a principal component of amino acid properties (PC1) strongly related to the hydrophobicity of the protein[6]. Repeating this same analysis after simulating library, replicate or DNA extraction bottlenecks in the original data results in lower correlations in all cases (Supplementary Figure 10, left column). Imposing both hard and soft minimum read count filtering as described above, the result is an increase in the correlation between measured fitness of amino acid substitutions and their corresponding predicted effects on PC1/hydrophobicity (Supplementary Figure 10, middle column).

Together this demonstrates that it is crucial to discard variants purely arising from sequencing errors to avoid systematic errors and shows how DiMSum can be used successfully to prioritize variants, minimize biases in downstream analyses and improve biological conclusions.

## Conclusions

We have developed a customizable pipeline – DiMSum – that provides a complete solution for the analysis of DMS data. DiMSum is easy to run, can handle a wide variety of different library designs, provides detailed reporting and produces fitness and error estimates form raw DNA sequencing data in a matter of hours. Importantly, DiMSum’s interpretable error model is able to identify and account for measurement errors in fitness scores resulting from random variability in DMS workflows and additionally provides the user with diagnostics to identify and deal with common causes of systematic errors. We have also shown that the DiMSum error model provides accurate error estimates across many published DMS datasets, outperforming previously used methods, and that diagnostic plots enable simple remedial steps to be taken that have the potential to dramatically improve the reliability of results from downstream analyses.

## Methods

### DiMSum software implementation

DiMSum is implemented as an R/Bioconda package and command-line tool compatible with Unix-like operating systems (see installation instructions: https://github.com/lehner-lab/DiMSum). The pipeline consists of five stages grouped into two modules that can be run independently: *WRAP* (DiMSum stages 1-3) processes raw FASTQ files generating a table of variant counts and *STEAM* (DiMSum stages 4-5) analyses variant counts generating variant fitness and error estimates. *WRAP* requires common software tools for biological sequence analysis (FastQC[39], cutadapt[40], VSEARCH[41] and starcode[58]) whereas *STEAM* has no external binary dependencies other than Pandoc. A detailed R markdown report including summary statistics, diagnostic plots and analysis tips is automatically generated. DiMSum takes advantage of multi-core computing if available. Further details and installation instructions are available on GitHub: https://github.com/lehner-lab/DiMSum.

### DiMSum data preprocessing

FastQ files from paired-end sequencing of the TDP-43 290-331 library[6] were processed with DiMSum v1.1.3 using default parameters with minor adjustments. First, 5, constant regions were trimmed in an error tolerant manner (‘cutadaptErrorRate’ = 0.2). Read pairs were aligned and those that contained base calls with posterior Phred scores (posterior score takes both Phred scores of aligned bases into account) below 30 were discarded (‘vsearchMinQual’ = 30, ‘vsearchMaxee’ = 0.5). Finally, variants with greater than two amino acid mutations were removed (‘maxSubstitutions’ = 2). One out of four input replicates (and all associated output samples) was discarded (‘retainedReplicates’ = 1,3,4) from all results shown in main text figures because the shape of its fitness distribution significantly differed from those of three other replicate experiments (see Supplementary Figure 4b,c). Note that Supplementary Figures 1-6 show DiMSum summary report plots when using all four replicates.

Simulated bottlenecked datasets were similarly processed with DiMSum v1.1.3 using hard, soft and no filtering. For datasets with library and replicate bottlenecks, filtering was performed on the Input samples only (‘fitnessMinInputCountAll’ = 10 for hard threshold or ‘fitnessMinInputCountAny’ = 10 for soft threshold), whereas for datasets with DNA extraction bottlenecks, Output samples were additionally filtered (‘fitnessMinOutputCountAll’ = 10 for hard or ‘fitnessMinOutputCountAny’ = 10 for soft threshold).

DMS datasets for leave-one-out cross validation were processed with DiMSum v1.1.3 except the data for Protein G B1 domain (GB1[5]) whose variant counts were obtained from Otwinoski 2019[59]. tRNA datasets[50] obtained from SRA (SRP134087) were analysed using DiMSum with default parameters except fitnessMinInputCountAll = 2000 and fitnessMinOutputCountAll = 200 to remove flaps likely due to DNA extraction bottlenecks, resulting in an average number of 2400 variants that could be analysed per selection experiment. The use of soft thresholds would result in an average increase in variant counts of 200% across the four selection experiments. For datasets with only one input sample (GB1 and tRNA), we replicated the input sample to create as many matched input-output samples as necessary for the error model analysis. All experimental design files and bash scripts with command-line options required for running DiMSum on the above datasets are available on GitHub (https://github.com/lehner-lab/dimsumms).

### DiMSum fitness estimation and error modelling

DiMSum calculates fitness scores of each variant *i* in each replicate *r* as the natural logarithm of the ratio between output read counts 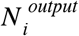 and input read counts 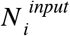 relative to the wild-type variant *wt*

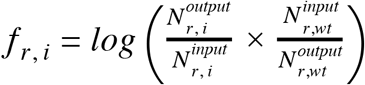

Optionally, DiMSum applies a scale and shift procedure to minimize linear differences in fitness scores between replicates. This is done by fitting a slope and an offset parameter to each replicate’s fitness scores in order to minimize the sum of squared deviations between variants, replicate fitness scores and their respective averages. Moreover, it is ensured that wild-type variants have an average fitness score of 0 across replicates.

Measurement error of fitness scores is modeled based on Poissonian statistics from sequencing counts of the variant 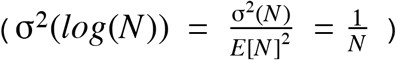, with multiplicative (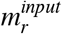 for input sample and 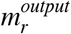 for output sample) and additive (*a_r_*) modifier terms that are common to all variants, but specific to each replicate experiment performed, as

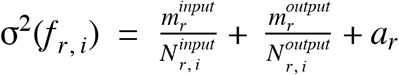

Note that we omit inclusion of error terms arising from the wild-type normalization, as these error terms are typically small due to high wild-type counts.

The error model is fit to a high confidence set of variants (variants that have enough sequencing reads in the input samples to display the full range of fitness scores and for which at least one sequencing read has been observed in all output samples, see Supplementary Figure 4a). The error model parameters are estimated by sharing information across all variants, that is by minimizing the sum over all variants’ squared deviation between the average error model prediction across replicates and the observed variance of fitness scores across replicates:

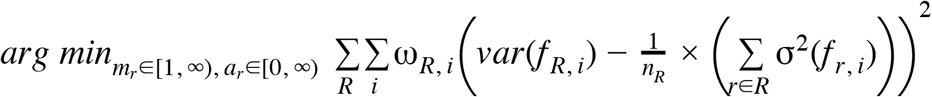

In order to reliably estimate replicate-specific additive error terms *a_r_*, the error model fit is performed not only with variances/error model predictions across all replicates of the DMS experiment, but across all possible subsets of replicates *R* of size at least two simultaneously (e.g. for three replicates, *R* ∈ ({1,2,3},{1,2},{1,3},{2,3}) and *n_R_* = {3,2,2,2}). This is done because estimation of additive error terms depends mostly on high count variants (which have little to no error contribution from sequencing counts) and the error model cannot distinguish how much additive variability was contributed by any one replicate unless further constrained (by subsets of lower order combinations). However, this means that if only two replicates of the DMS experiment have been performed, the error model tends to split additive error contributions equally between replicates for lack of more information, i.e. additive error terms cannot be used as diagnostic.

Moreover, squared deviations between variance and average error model predictions per variant and replicate subset are weighted (*ω_R,i_*) according to three factors. First, the number of replicates in the replicate subset R, to account for differential uncertainty in empirical variance estimates. Second, the inverse of the average count-based error according to Poissonian statistics, to minimize relative, not absolute, deviations between variance of fitness scores and respective error model estimates. Third, a term re-weighting all variants with the same number of mutations according to 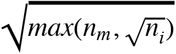, where *n_m_* is the number of variants with that number of mutations and *n_i_* is the overall number of variants in the high-confidence variant pool. This is done to place more weight on the typically fewer lower order mutant variants (e.g. single mutants) and therefore to improve estimates of additive error terms.

The error model is fit 100 times on bootstrapped data. For each bootstrap, at maximum 10,000 variants are drawn with replacement from the high-confidence variant pool. The average parameters across bootstraps are used to calculate measurement error estimates. The error estimates are then used to merge fitness scores across replicates by weighted-averaging

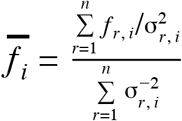

The corresponding error of these merged fitness scores is calculated as

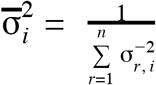

DiMSum reports merged fitness scores and associated errors for all variants that have been observed in at least one experimental replicate (actual merging is performed for variants observed in two or more replicates; for variants only observed in one experimental replicate, merged fitness scores and error are simply those computed for this one replicate).

DiMSum diagnoses consistency of the error model with the data by estimating how well it describes fitness score differences between replicates. If all error sources have been accounted for and model parameters accurately attribute error contributions to the different replicates, the predicted error magnitude should match the randomly arising differences in fitness scores between replicates of the same experiment, which we find to be normally distributed across all DMS datasets investigated, i.e.

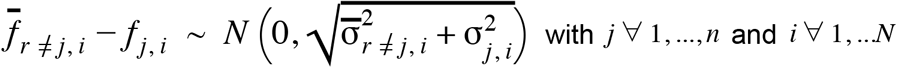

We thus calculate a z-score of the fitness score differences replicates as

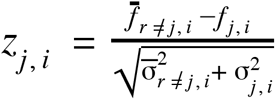
that should follow a Normal distribution centered on zero and with unit standard deviation. DiMSum outputs quantile-quantile plots of *Z_j, i_* as well as its corresponding mean and standard deviations, and the P-value distribution from a two-sided z-test (Supplementary Figure 6e,f). Mean values of *Z_j, i_* different from zero suggest that fitness score estimates are biased, which suggests the presence of systematic errors not accounted for by the ‘scale and shift’ normalization procedure and the error model. A standard deviation different from one suggests that the error has been overestimated (s.d. < 1) or underestimated (s.d. > 1).

### Error model validation and benchmarking

#### Artificial increases in multiplicative and additive error terms in Supplementary Figure 7

In order to show that the error model can accurately capture multiplicative and additive error sources, we performed a manipulation of the TDP-43 290-331 library (using only replicates 1, 3 and 4) in which we artificially increased multiplicative input terms (by multiplying input count reads by factor 3 or 10) or additive error terms (by adding values of 0.3 or 1 to normalized fitness scores immediately before error model fitting) for replicate 1.

#### Leave-one-out cross validation

To benchmark the error model and compare it against alternative approaches to quantify measurement error, we performed leave-one-out cross validation on published DMS datasets. In contrast to the error model benchmarking performed as a diagnostic output from the DiMSum pipeline (see above), we trained the error models on all but one replicate of a dataset in turn. These error models were then used to calculate a z-score of the fitness score differences between the unseen replicate and the average over the training replicates (*r* ≠ *j*) as

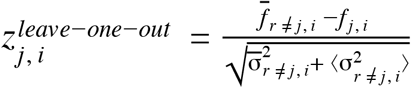
where 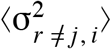 is the prediction of error in the test replicate *j* using the error model parameters of the training replicates.

For the DiMSum error model, this is 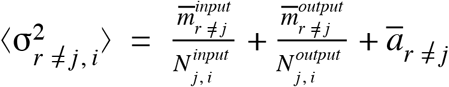 with 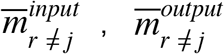 and 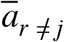 as the averages of replicate-specific error model terms over the training replicates.

In Figure 3a,c 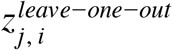 is compared against a Normal distribution, with the expectation that they should match if the error magnitude is correctly predicted. Figure 3b,d displays the P values from a two-sided z-test using 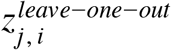, with the expectation that P values should be uniformly distributed, because there should not be significant differences of fitness scores between replicate experiments. Finally, to analyse 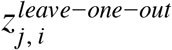 systematically across many datasets, we calculated the inverse of its standard deviation for each dataset (Table 2, Figure 3e, Supplementary Figure 8). I.e. 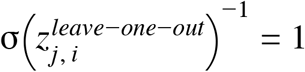 if the error model has correctly predicted the magnitude of measurement errors, 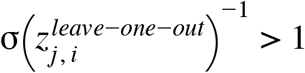 if errors have been over-estimated or 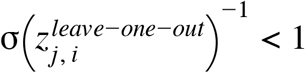 if errors have been under-estimated.

**Table 2:**
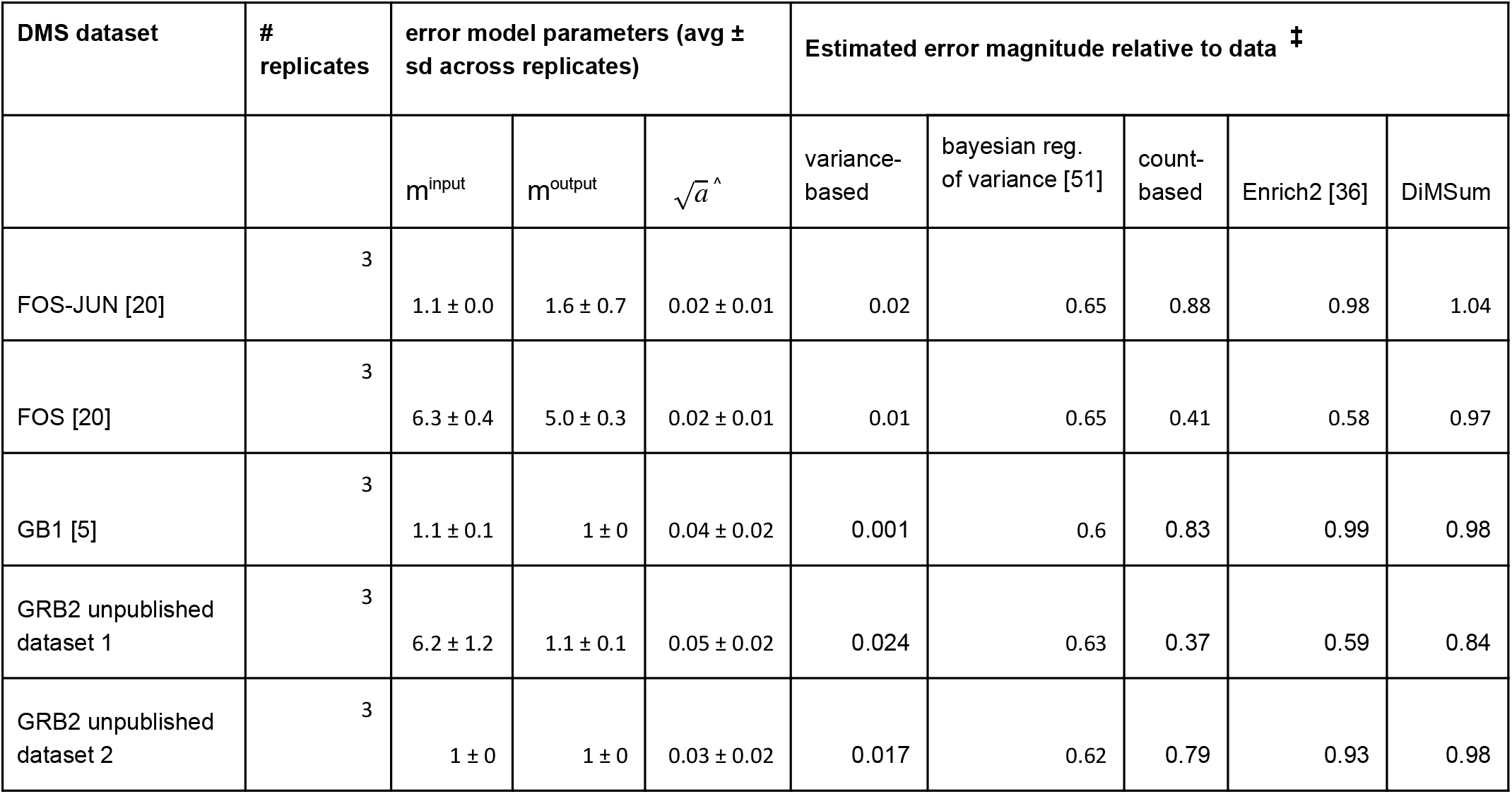
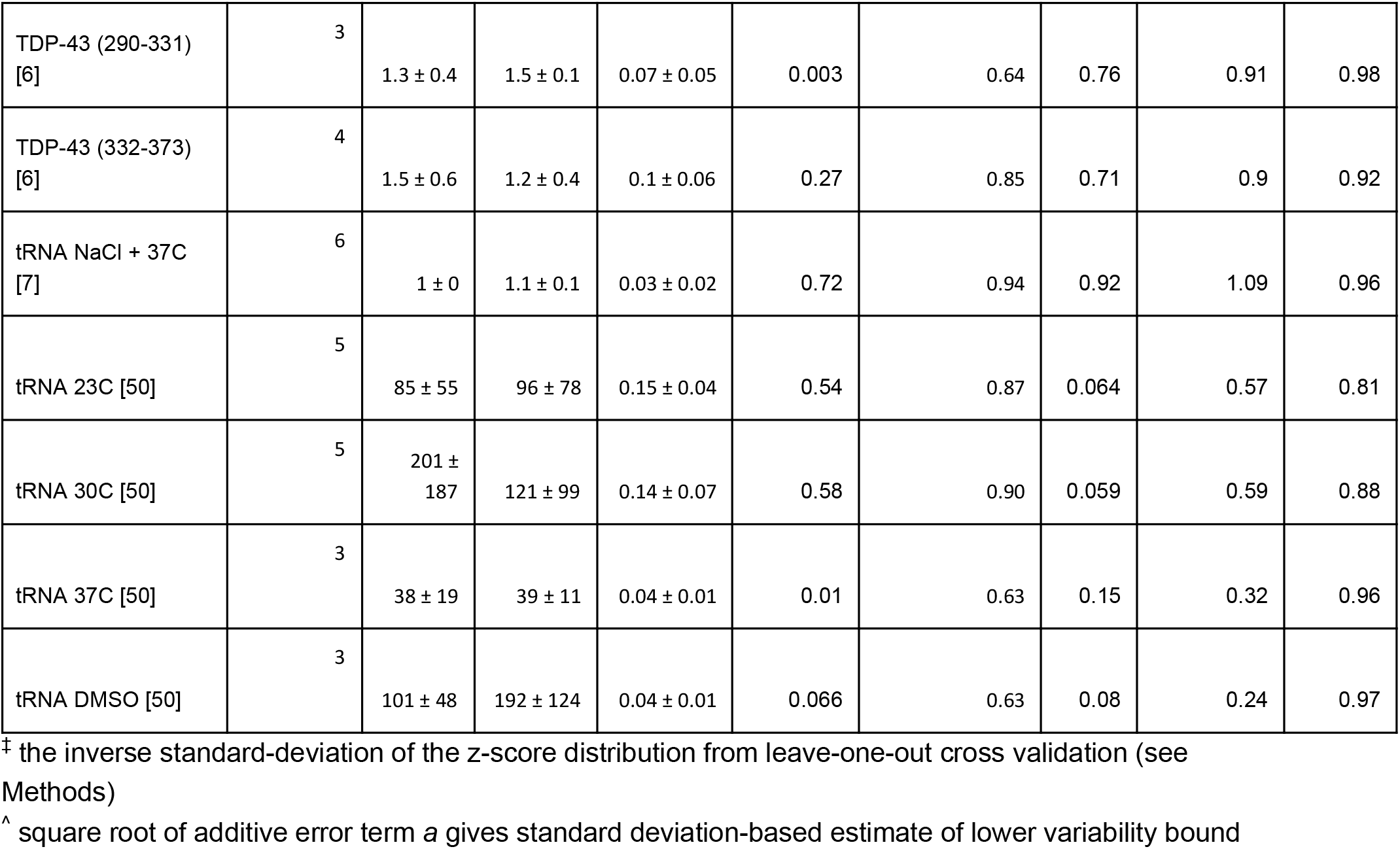
DiMSum error model parameters and error model performance in leave-one-out cross validation across twelve DMS datasets

### Alternative error models

We compared the DiMSum error model to four alternative error models.

First, a ‘variance-based’ error model, where the error of fitness scores for each variant is calculated from the empirical variance of fitness scores between replicates, i.e. with the measurement error of fitness scores merged across replicates as 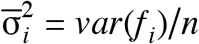.

Second, an error model using a Bayesian regularization of the empirical variance, as introduced by Weile et al. [51]. Here the empirical variance of each variant’s fitness scores between replicates is regularized with a prior, which is a regression of the empirical variance on input sequencing counts and fitness scores. Here the measurement error of fitness scores merged across replicates is 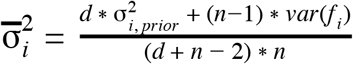, with *d* as the degrees of freedom of the regression, 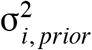 the prior estimate of the variance for variant *i and n the number of replicate experiments*. For the variance-based error models, the measurement error for the unseen test replicate was estimated as the average of individual training replicates. The z-scores for the variance-based error model in the leave-one-out cross validation were thus calculated as 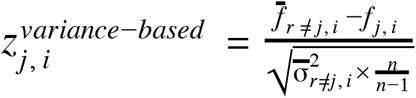

Third, a minimal ‘count-based’ error model, where the error of fitness scores is estimated from sequencing read counts in Input and Output samples under the assumption that sequencing counts follow a Poisson distribution, i.e. as for the DiMSum error model but without multiplicative or additive terms.

Fourth, the Enrich2 error model by Weile et al. [36], which is based on sequencing counts but modified with variant-specific correction terms (‘random-effect model’). Here error estimates are calculated from input and output sequencing read counts under Poisson assumptions, but with an additional variant-specific random-effect term. This term corrects error estimates if the observed variability of fitness scores across replicates is larger than the estimated count-based error alone. That is, if 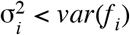, the random-effect term 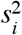 is estimated greater 0 such that 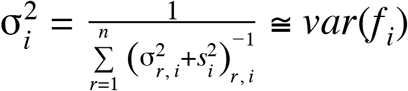, i.e. the error estimate becomes equivalent to that of the variance-based error model described above. To calculate z-scores in the leave-one-out cross validation for the Enrich2 error model, we estimated random-effect terms across the training replicates, and then also used them to modify the count-based error estimate of the test replicate, i.e. 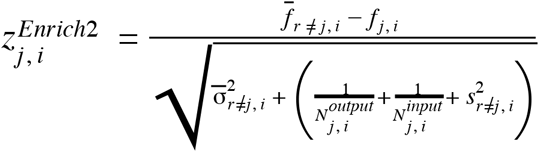.

### Multiplicative errors from PCR amplification

Kowalsky et al.[54] previously reported increased variability in sequencing read counts due to PCR amplification protocols (see Table S3 of Kowalsky et al.[54]). The raw sequencing data for the three PCR amplification protocols tested was obtained from the authors. Paired-end reads were merged with USEARCH[60] using the *usearch -fastq_mergepairs* command with a minimum per base posterior Qscore of 20 and reads for unique variants were counted using the *usearch -fastx_uniques* command. To allow estimation of multiplicative and additive error terms, we treated the sequencing data from each PCR amplification protocol as replicate experiments. Variant fitness scores were calculated as the natural logarithm of read count frequency (read counts divided by total number of reads in each replicate). Error of fitness scores was calculated as the inverse of variant read counts. DiMSum error model was adjusted to only fit one multiplicative error term and the additive error term per replicate. Additive errors were small compared to variability observed. Multiplicative error terms were 1.9 ± 0.4 for Method A (using one amplification cycle with all primers at once), 1.4 ± 0.1 for Method B (two amplification cycles interspersed with a ExoI degradation step) and 6.4 ± 0.6 for Method C (two amplification cycles).

### Simulated bottlenecks in a previously published DMS dataset

We used a DiMSum processed DMS dataset from Bolognesi and Faure et al.[6] (290-331 library) to simulate the effects of various experimental bottlenecks.

#### Simulating a Library Bottleneck

A library bottleneck of size α = 0.03 (meaning that only 3% of molecules pass through the bottleneck) was simulated based on the observed average frequencies of variants in the input samples. A bottleneck factor 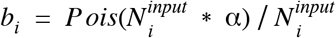 was calculated to capture the subsequent changes in read count frequencies that would occur during such a bottleneck. For variants with high counts in input samples, the bottleneck factor will be close to α. However, for low count variants, the bottleneck factor will vary considerably. Some variants, especially variants with 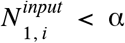, will not pass through the bottleneck, i.e. *b_i_* =0, while others may pass through the bottleneck even though there was only one molecule of that variant present in the pool, i.e. *b_i_* =1.

To simulate how read counts in sequencing samples (both input and output sequencing samples) change due to this bottleneck, we sampled *N* times from a multinomial distribution *Mult*(1, π_*r*_) where *N* is the total number of sequencing reads in the ‘original’ sample *s* and π_*s*_ is a vector of probabilities given by:

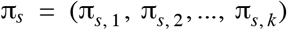

where *k* is the total number of different variant sequences, and π_*r, i*_ is the frequency of variant *i* in sample *s* after the library bottleneck (e.g. for replicate *1* output sample):

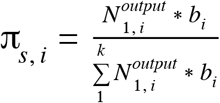

To simulate sequencing errors in the new modified data, we assumed that the probability that a given sequencing read is misidentified to be 0.02, based on a length of the mutated sequence of 126nt and a per base misread frequency of 0.0001[6], and that all errors involve WT molecules being misclassified as single mutants, or single mutant molecules being misidentified as double mutants, or double mutant molecules being misidentified as triple mutants, and triple mutant molecules being misidentified as quadruple mutants. The total number of triple mutant molecules that will be mis-identified as quadruple mutants is 0.02*N*, where *N* is the total number of triple mutant reads. Those counts were randomly subtracted from the triple mutant counts, and added to the counts of all the quadruple mutants. The total number of double mutant molecules mis-identified as triple mutants is 0.02*N*′, where *N*′’ is the total number of double mutant reads. Those counts were subtracted from the double mutant counts, and randomly distributed among all the triple mutants. This process was repeated to simulate single mutants being mis-identified as double mutants, and WT molecules being mis-identified as single mutants.

#### Simulating a Replicate Bottleneck

The procedure was similar to the library bottleneck procedure described above, but a bottleneck factor was calculated on each replicate input sample independently, allowing for different variants to be present in each replicate.

#### Simulating a DNA Extraction Bottleneck

The procedure was similar to the library/replicate bottleneck simulations, but a bottleneck factor was calculated for each sequencing sample independently

## Declarations

### Ethics approval and consent to participate

Not applicable.

### Consent for publication

Not applicable.

### Availability of data and materials

Raw datasets analysed during the current study (see Table 2) are available in the GEO repository:

https://www.ncbi.nlm.nih.gov/geo/query/acc.cgi?acc=GSE102901

https://www.ncbi.nlm.nih.gov/geo/query/acc.cgi?acc=GSE128165

https://www.ncbi.nlm.nih.gov/geo/query/acc.cgi?acc=GSE99418

https://www.ncbi.nlm.nih.gov/geo/query/acc.cgi?acc=GSE111508

All experimental design files and bash scripts with command-line options required for running DiMSum on the above datasets can be found on GitHub: https://github.com/lehner-lab/dimsumms. We also provide an R package with custom scripts to perform all downstream analyses, simulations, performance comparisons and reproduce all figures described here.

DiMSum source code, installation instructions, command-line options, common use cases, input file formats (example templates) and the DiMSum demo mode are available on GitHub: https://github.com/lehner-lab/DiMSum.

### Authors’ contributions

All authors jointly conceived the project. A.J.F. and J.M.S. developed the pipeline software and performed analyses of DMS datasets. J.M.S. developed the error model. P.B. conducted bottleneck simulations. A.J.F. and J.M.S. wrote the manuscript with input from B.L. and P.B.

## Acknowledgements

The authors thank members of the Lehner lab for feedback on the DiMSum pipeline and methods.

## Supplementary Figure legends

**Supplementary Figure 1**. **DiMSum pipeline report: Raw FastQ file quality control summary and constant region trimming statistics. a.** 10^th^ percentile (upper) and mean (lower) Phred quality scores shown for forward reads (Read 1) in all FastQ files (see legend). **b**. Similar to A except quality scores for reverse reads (Read 2) are shown. Vertical dashed lines separate variable from constant regions. **c.** Percentage of forward reads (Read 1) in which corresponding 5’ and/or 3’ constant regions were matched and trimmed (see legend), shown separately for each FastQ file. **d.** Similar to A except trimming statistics are shown for reverse reads (Read 2).

**Supplementary Figure 2**. **DiMSum pipeline report: Paired-end read alignment statistics and variant processing and mutation statistics. a.** Percentage of successfully aligned paired-end reads (“vsearch_aligned”) retained for downstream analysis. Remaining read pairs not matching user-specified criteria are discarded. **b**. Aligned read length distributions shown separately for all samples (see legend). **c.** Per-sample total counts (and percentages, **d**) of retained processed reads with 0,1,2 or >3 (3+) nucleotide substitutions (see legend), as well as those discarded due to “invalid barcodes” (in the case of barcoded libraries), “indel” mutations, mutated “internal constant regions”, non-intended mutations (“not permitted”), variants with more substitutions than desired (“too many”) and nonsynonymous variants that have synonymous substitutions in other codons (“mixed”). **e** and **f** are similar to a and b, but indicate total counts and per sample percentages of read amino acid mutation statistics respectively (shown if the target molecule was a coding sequence).

**Supplementary Figure 3**. **DiMSum pipeline report: Marginal input variant count distributions and inter-sample variant count diagnostic plot. a.** Input sample variant count distributions for nucleotide substitution variants with hamming distances to the wild type sequence of up to 6. Wild-type counts are indicated by the black dashed line. Expected “fictional” variant count frequencies are indicated by red and brown dashed lines for single and double nucleotide substitution variants respectively. Bimodal distributions and unimodal distributions not surpassing indicated thresholds are indicative of variants originating from sequencing errors due to a library bottleneck (see Figure 4a). **b**. Scatterplot matrix depicting correlation between all Input and Output sample variant counts. Matrix cells in the upper triangle show Pearson correlation coefficients. Matrix cells in the lower triangle show hexagonal heatmaps, where each hexagon is shaded by corresponding two dimensional bin counts. Matrix diagonal cells indicate count densities. Distinct variant populations or ‘flaps’ – subsets of variants that appear at high counts in one replicate but at low counts in another – not resulting from applied selection are indicative of replicate or DNA extraction bottlenecks (see Figure 4b).

**Supplementary Figure 4**. **DiMSum pipeline report: Input read threshold for full fitness range and fitness normalisation. a.** Two dimensional hexagonal heatmaps showing replicate fitness scores versus Input variant counts. The vertical dashed line indicates the minimum Input count threshold used for retaining variants (covering the full fitness range) for subsequent error model fitting. **b**. Replicate fitness distributions before (and after, **c**) ‘scale and shift’ inter-replicate normalisation. Deviations in distribution shape (e.g. replicate 2) indicate systematic errors between replicates; affected replicates might have to be excluded from error model fitting and downstream analyses.

**Supplementary Figure 5**. **DiMSum pipeline report: Inter-sample variant fitness diagnostic plot.** Two dimensional hexagonal heatmaps showing replicate fitness score correlations (cells in the lower matrix triangle), corresponding Pearson correlation (upper triangle) and marginal densities (matrix diagonal). Only variants retained during error model fitting are depicted.

**Supplementary Figure 6**. **DiMSum pipeline report: Fitness error model estimates. a**. and **b**. Multiplicative (panel a) and additive (panel b) error terms estimated by the error model on TDP-43 290-331 dataset. Dots give mean parameters, error bars 90% confidence intervals. **c**. Empirical variance of replicate fitness scores as a function of error estimates purely based on sequencing counts under Poisson assumptions. Empirical variance (blue dots show average variance in equally-spaced bins, error bars indicate avg. variance(1 + 2 / #variants per bin)) is over-dispersed compared to baseline expectation of variance being described by a Poisson distribution. The bimodality of the count-based error distribution results from relatively few single nucleotide mutants with high counts (thus low count-based error) and many double nucleotide mutants with low counts (thus higher count-based error). The DiMSum error model (red line) accurately captures deviations of the empirical variance from Poisson expectation (black dashed line). **d**. Cyan and magenta lines indicate multiplicative error term contributions corresponding to Input and Output samples respectively. The horizontal green line indicates the additive error term contribution. The red line indicates the full DiMSum error model. **e, f**. DiMSum outputs quantile-quantile plots of z-scores of the differences between replicate fitness scores as a visual check whether the error model has properly estimated error magnitudes. Z-scores are shown for individual replicates (the indicated replicate is the ‘test’ replicate) and aggregated across all replicates (grey, as shown in Figure 3 for different error models). Shown here is the difference between DiMSum error models when replicate 2 is included in the analysis (panel **e**) or excluded from the analysis (panel **f**). In the first analysis, replicate two shows biased fitness scores, while the latter analysis the three remaining replicates have unbiased fitness scores with the correct error magnitude.

**Supplementary Figure 7**. **DiMSum error model estimates multiplicative and additive error sources in fitness scores. a**. and **b**. Multiplicative (panel **a**) and additive (panel **b**) error terms estimated by the error model on TDP-43 290-331 dataset after multiplicative manipulation (middle and right panels) of the original data (left panels). **c**. and **d**. Multiplicative (panel **c**) and additive (panel **d**) error terms after additive manipulation (middle and right panels) of the original data (left panels). The DiMSum error model captures multiplicative (panel **a**) and additive (panel **d**) manipulations of the original dataset, with minor effects on non-manipulated parameters. Of note, manipulating replicate 1 input multiplicative error results in minor lowering of replicate 1 output multiplicative error, possibly due to slight overlap in variants that are similarly affected by both error terms. Additionally, at very high addition of replicate 1 additive error, multiplicative errors are affected. This is likely because at a magnitude of 1 (s.d.) the additive error dominates the overall error (and exceeds by far any additive error term observed in published DMS datasets), masking error contributions by other error sources and thus complicate their reliable estimation. Dots give mean parameters, error bars 90% confidence intervals.

**Supplementary Figure 8**. **Leave-one-out cross validation of different error models on twelve DMS datasets**. Leave-one-out cross validation of different error models on twelve DMS datasets stratified by single and double mutants and also showing performance of partial DiMSum error models. Color gives an estimate of the over-dispersion in the datasets (estimated as average over all multiplicative error terms of the DiMSum error model; yellow – litte, blue – a lot) to show dependency of the performance of count-based error models on over-dispersion. Point shape indicates the number of replicates in the dataset to show dependency of the performance of empirical variance based error models on the number of replicates. Single or double mutants are on nucleotide level (tRNA datasets) or amino acid level (all other datasets). ‘DiMSum additive error’ indicates DiMSum error model using only additive error terms (setting multiplicative terms to 1). ‘DiMSum multiplicative error’ indicates DiMSum error model using only multiplicative error terms (setting additive terms to 0).

**Supplementary Figure 9**. Scatterplots of Input and Output counts of previously published DMS experiments[20,50]. Only variants with two nucleotide substitutions are shown. For FOS and FOS-JUN datasets, counts of single AA variants with two nucleotide substitutions in the same codon are shown. The black arrows indicate populations of spurious double nucleotide variants originating from sequencing errors.

**Supplementary Figure 10.** Effects of bottlenecks and variant filtering on fitness estimates and biological conclusions. Relative fitness of single and double amino acid variants as a function of hydrophobicity changes induced by mutation. Results are shown separately for the original data (no bottlenecks) as well as after simulating library, replicate and DNA extraction bottlenecks (3%). The middle column indicates the results after imposing a hard variant count threshold of 10 reads. Relative fitness was calculated from changes of output to input frequencies with respect to the WT sequence. The rightmost column indicates the results after imposing a soft variant count threshold of 10 reads (the original data was not filtered). For the DNA extraction bottleneck, read count thresholds were similarly applied to output samples.

