## Supplementary figures for "DiMSum: an error model and pipeline for analyzing deep mutational scanning data and diagnosing common experimental pathologies"

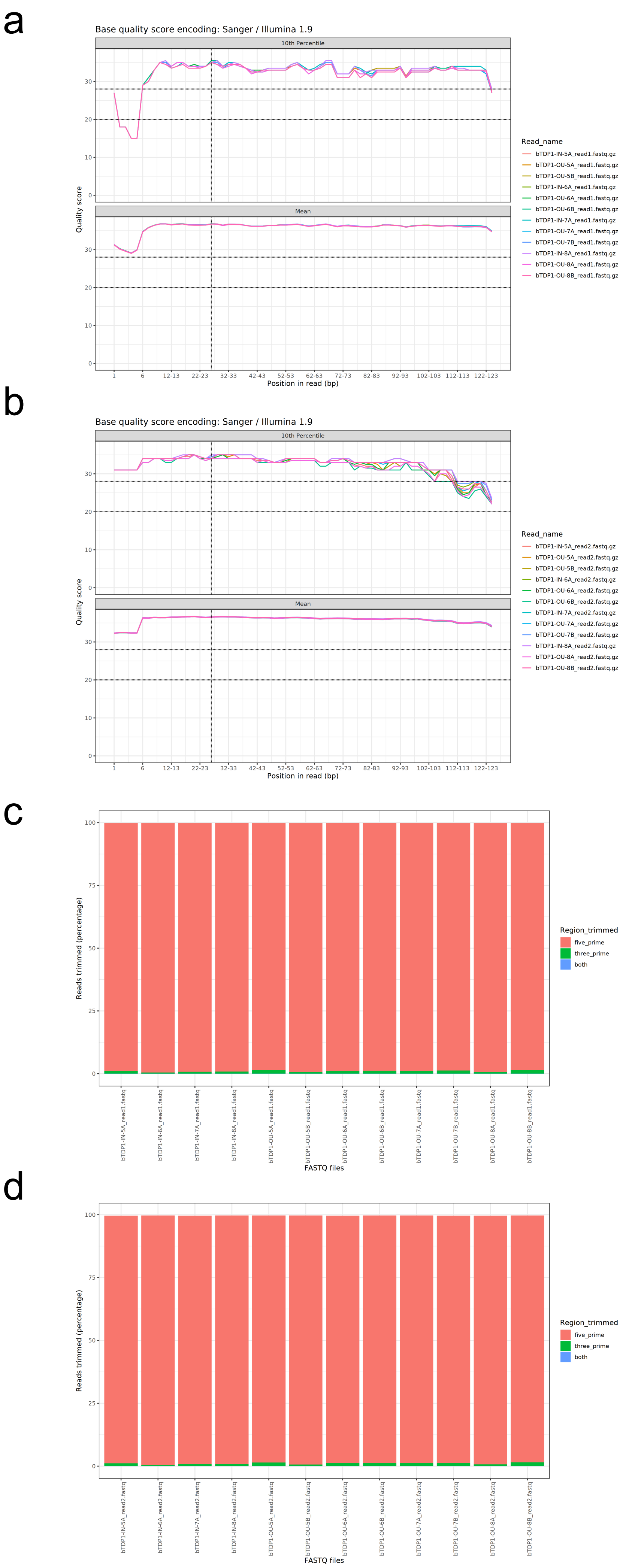


Supplementary Figure 1

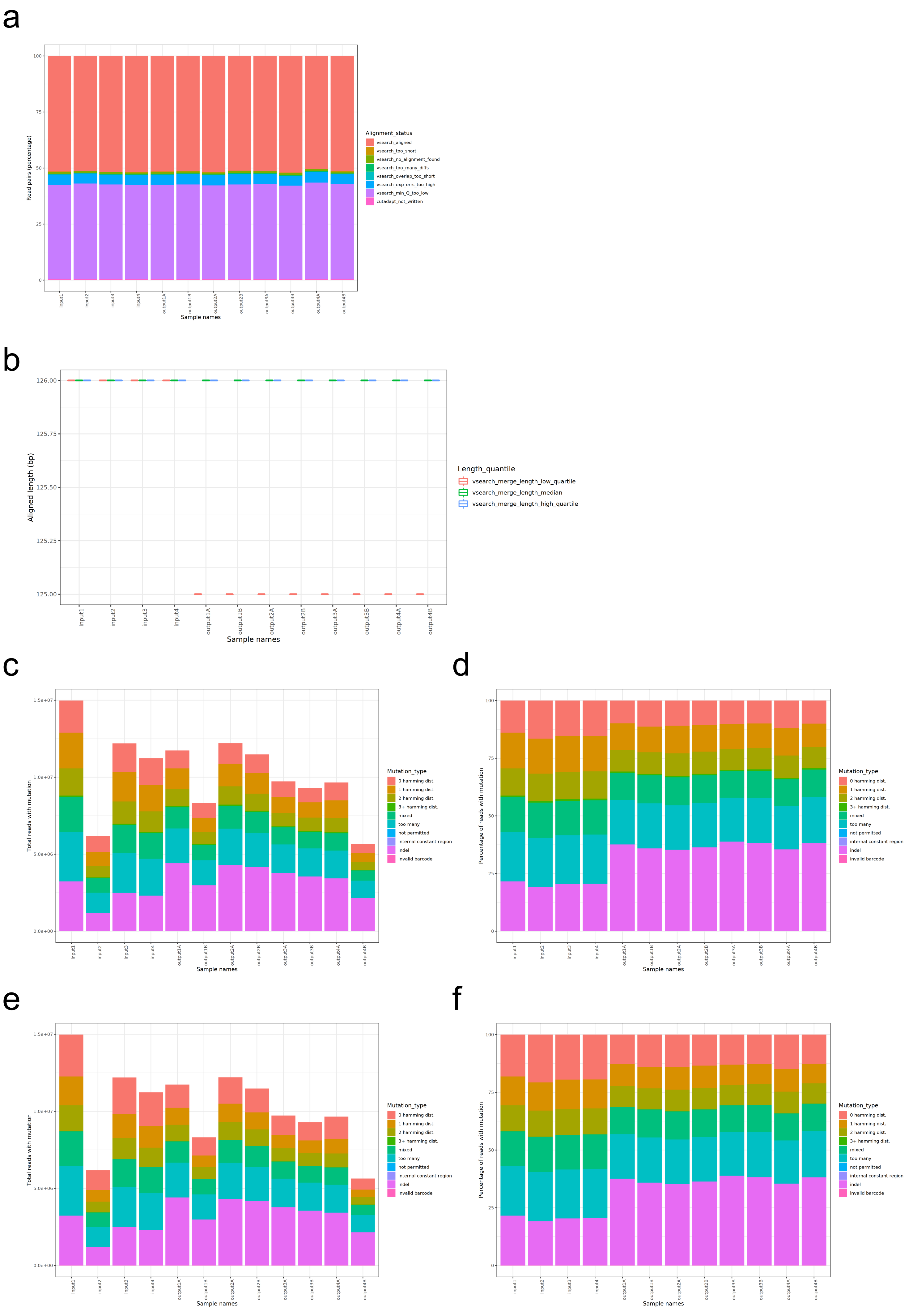


Supplementary Figure 2

a

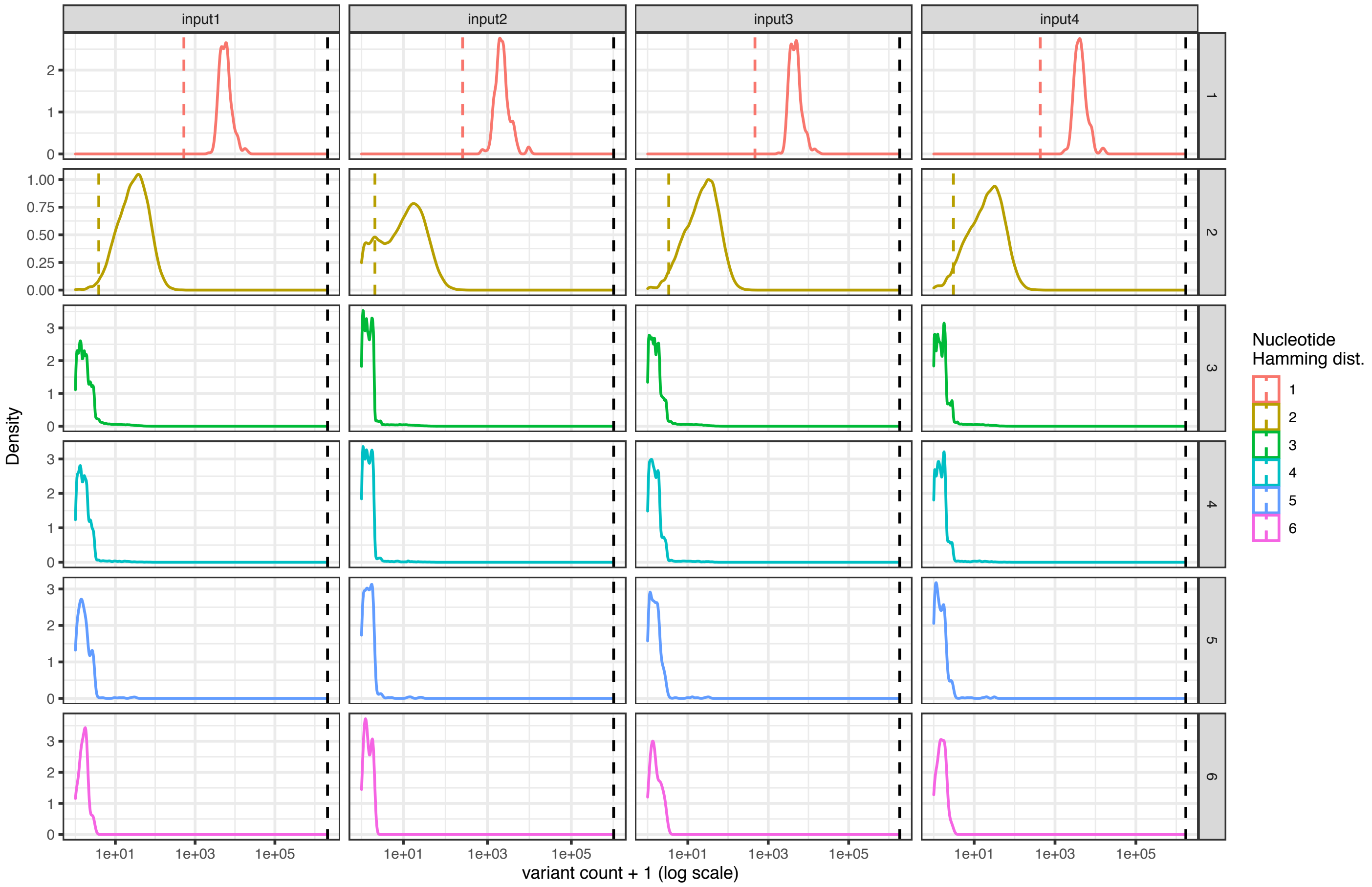


b

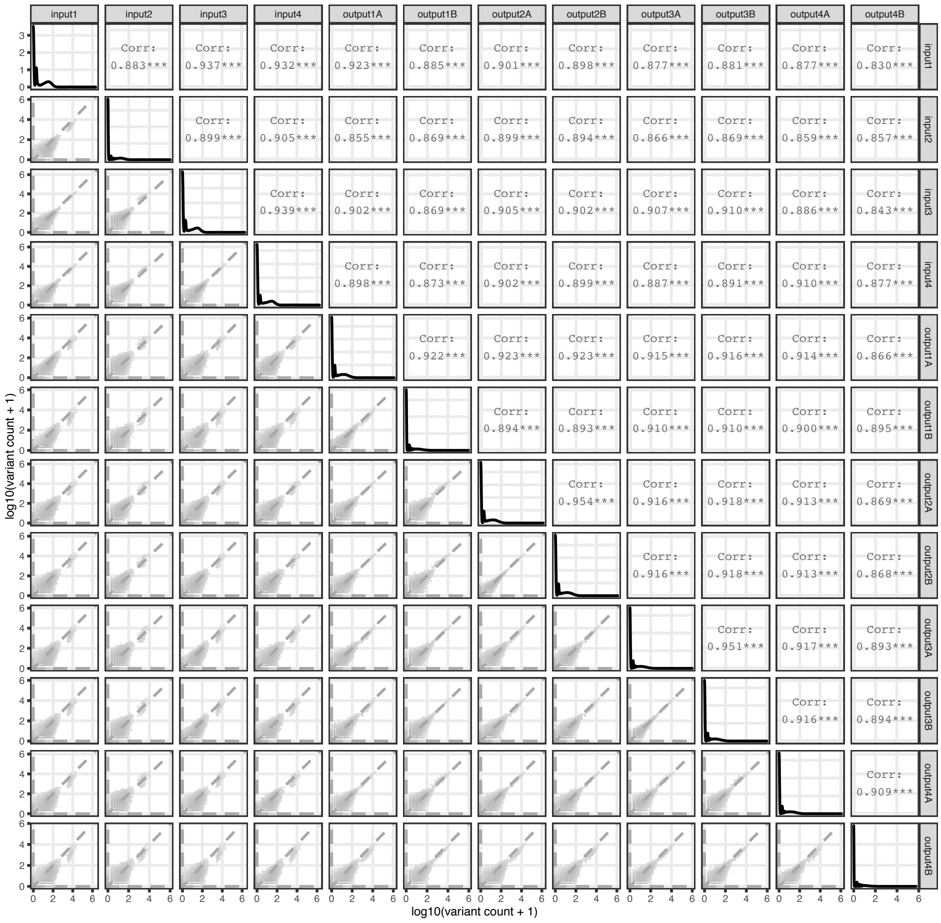


### Supplementary Figure 3

a

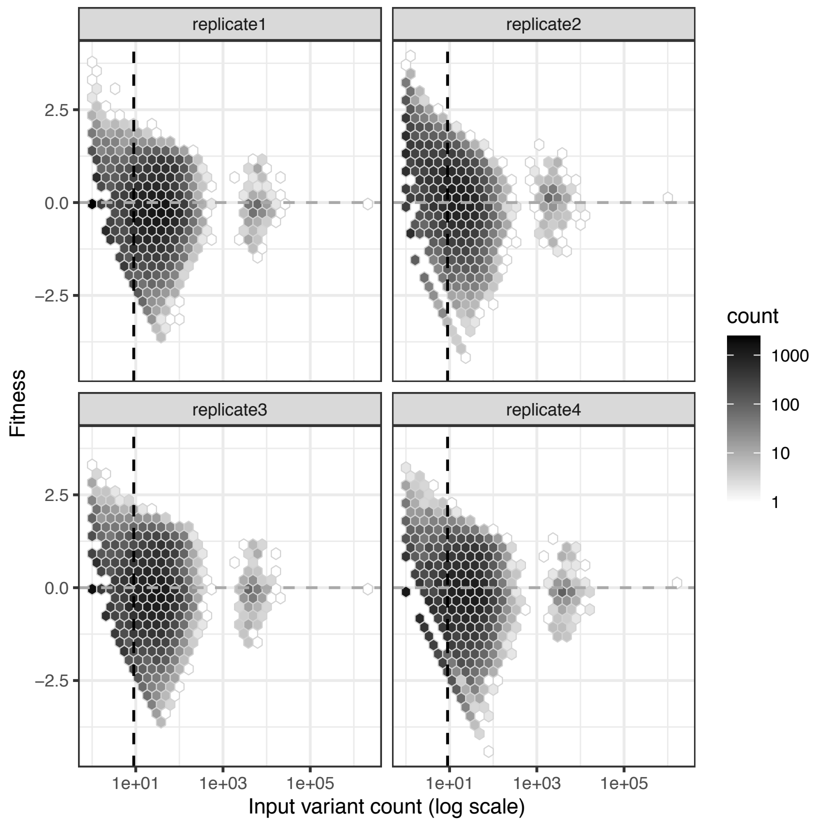


b

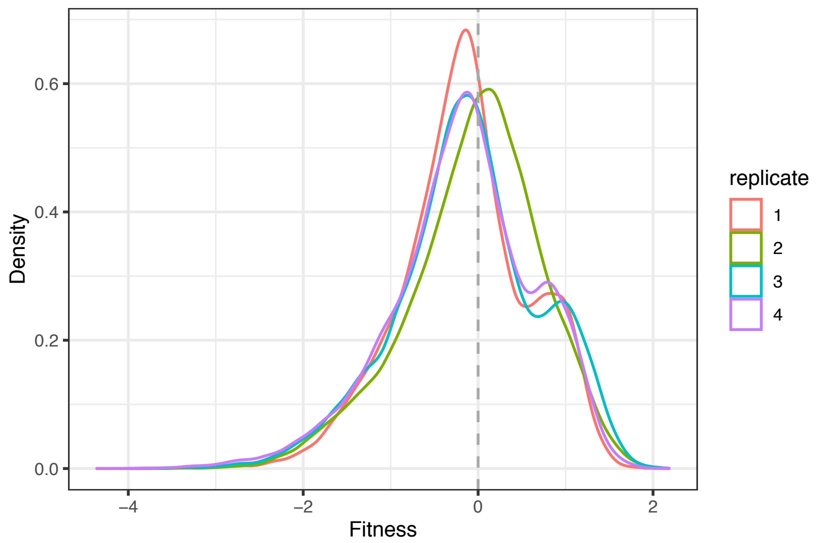


c

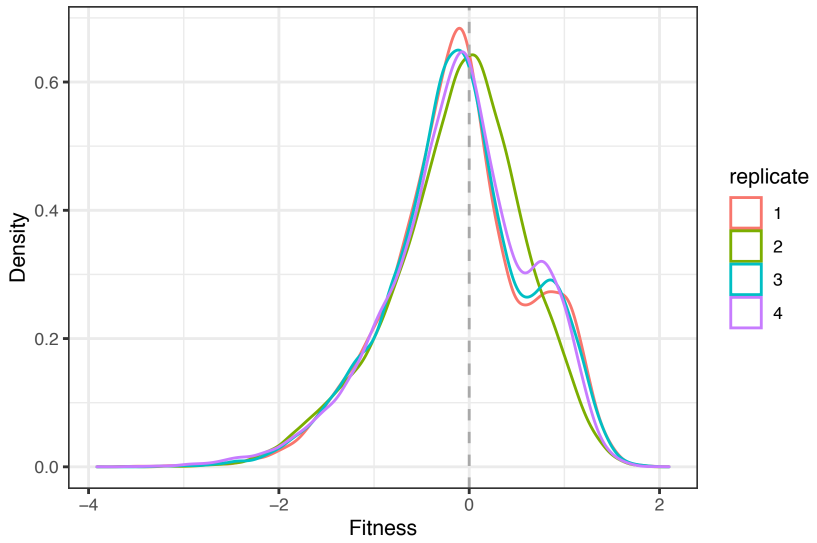


### Supplementary Figure 4

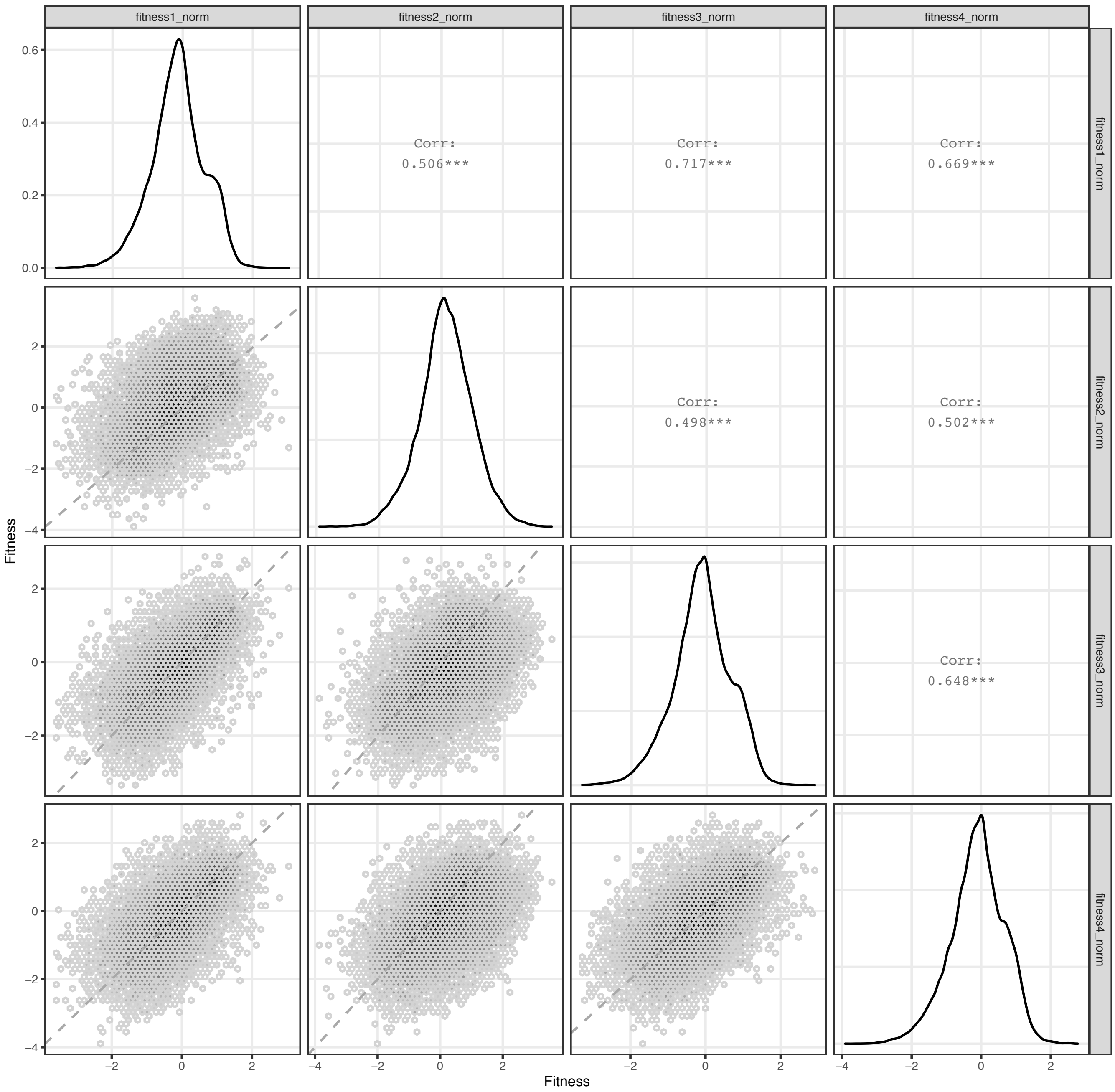


Supplementary Figure 5

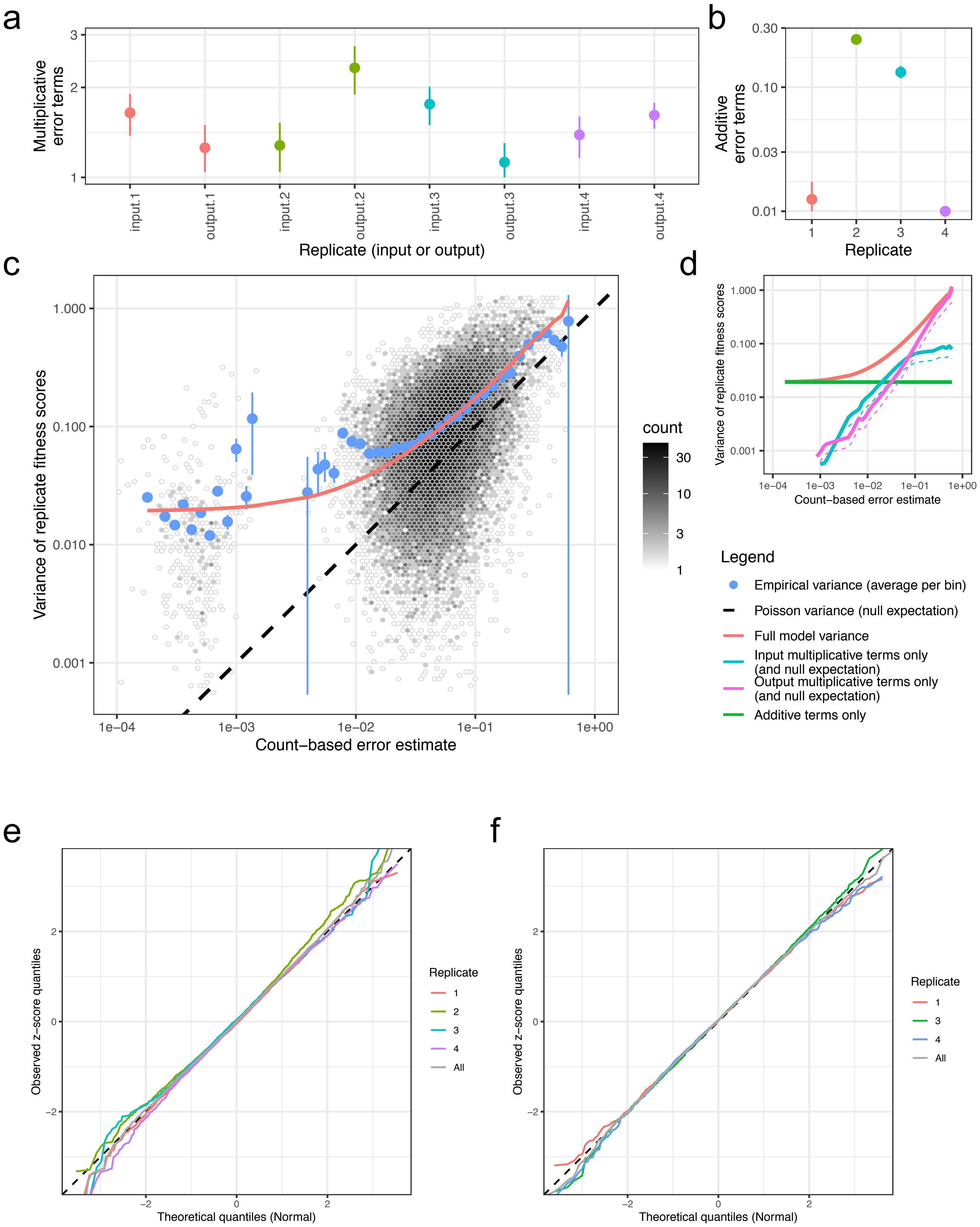


### Supplementary Figure 6

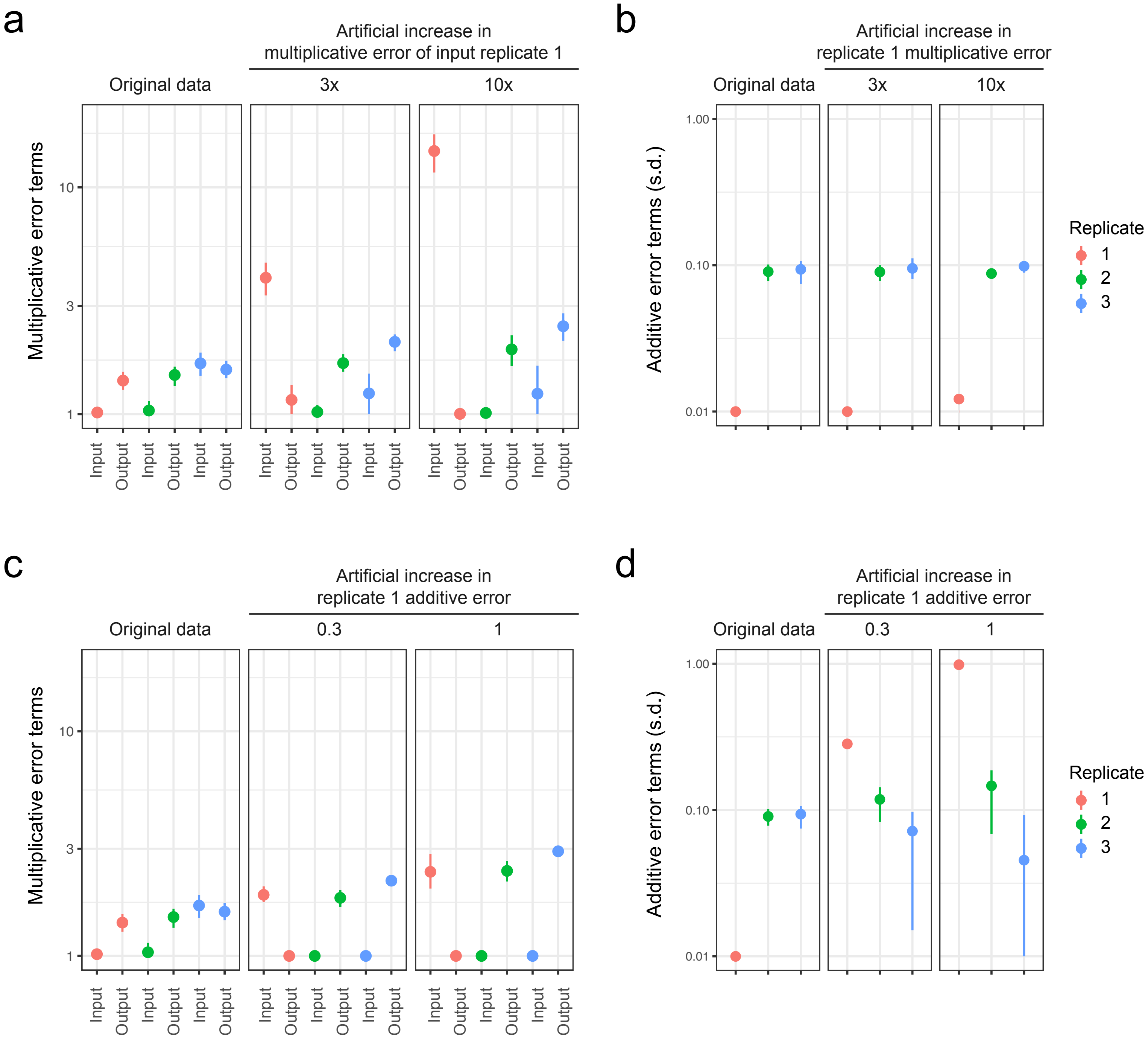


Supplementary Figure 7

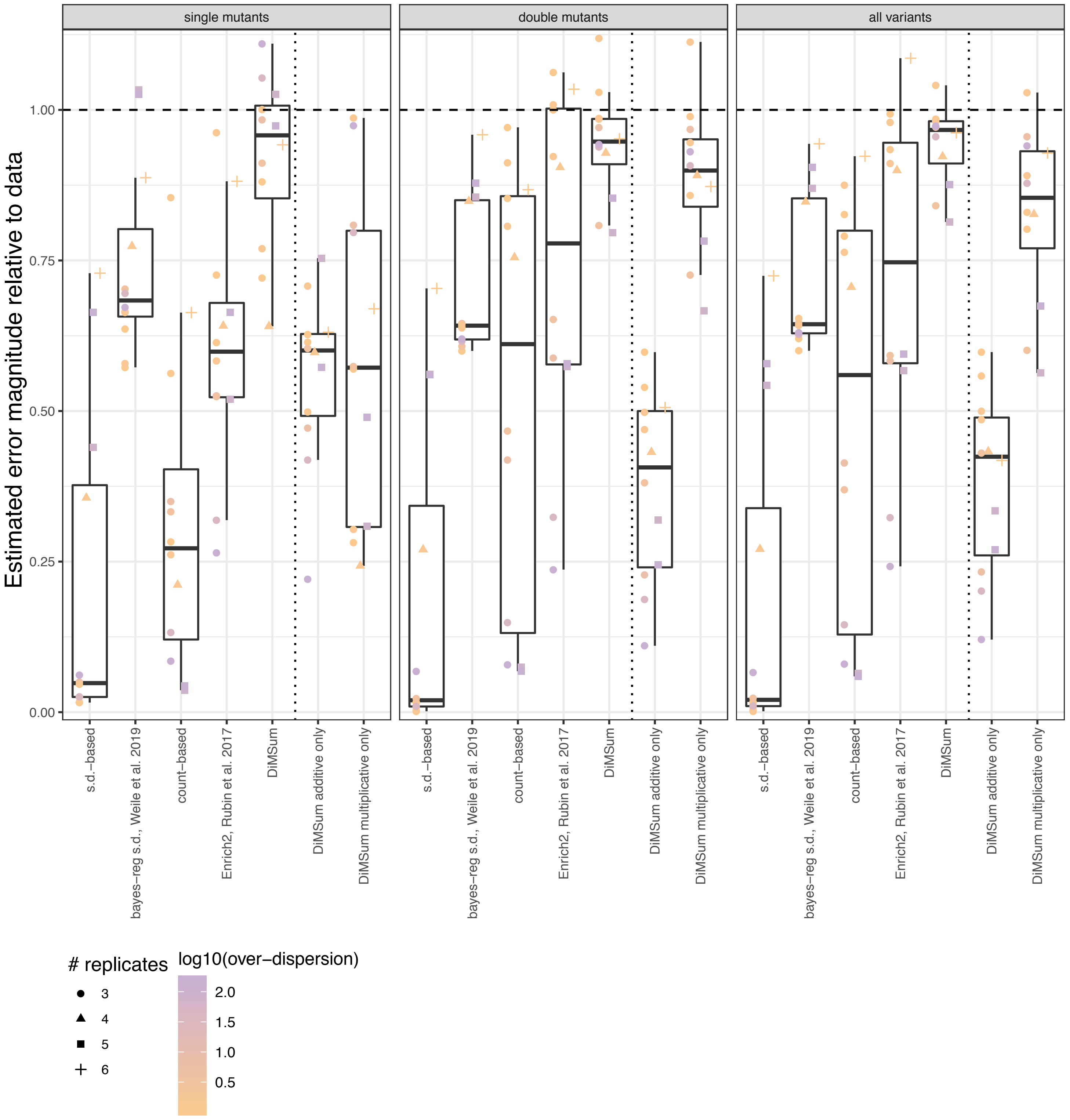


Supplementary Figure 8

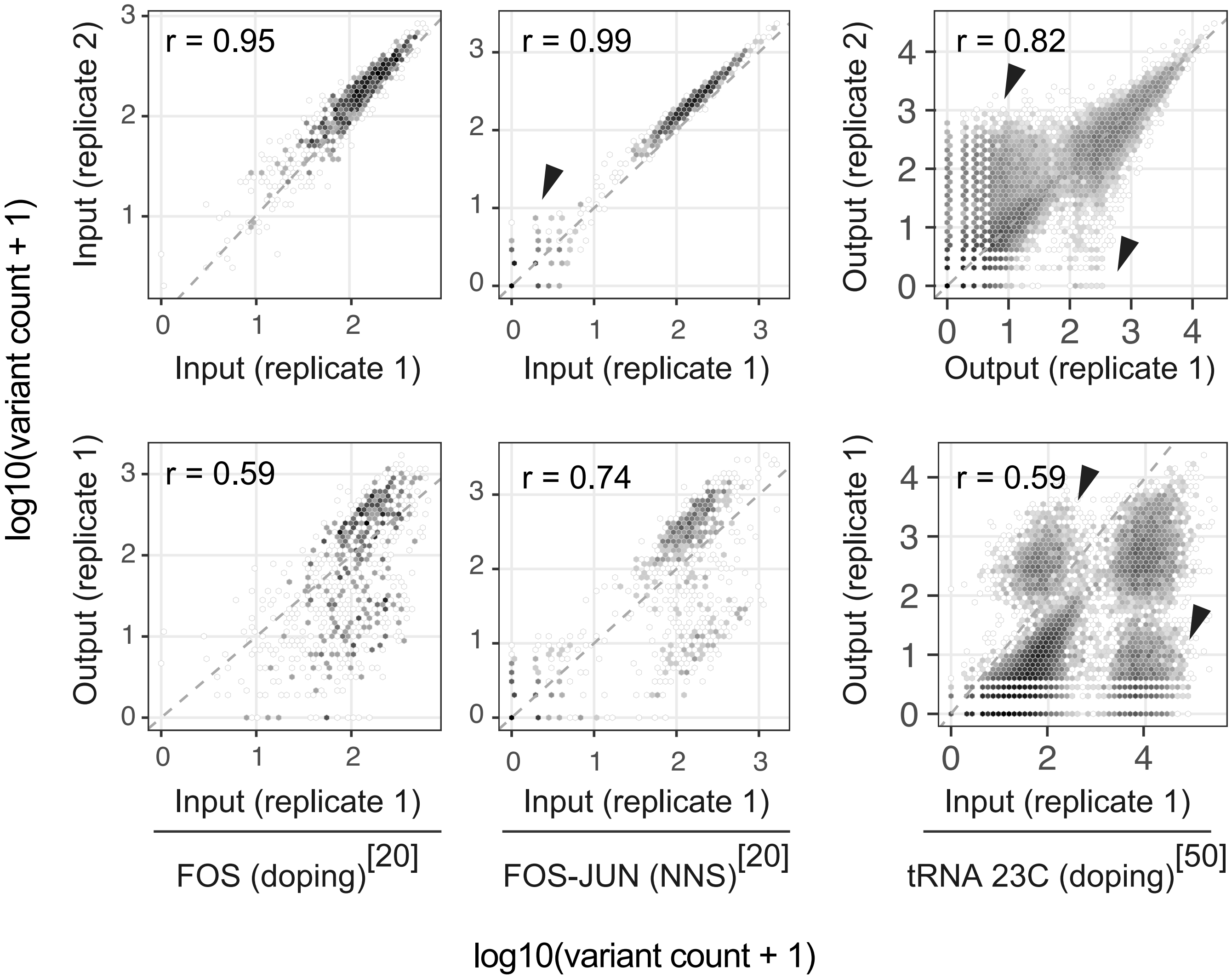


Supplementary Figure 9

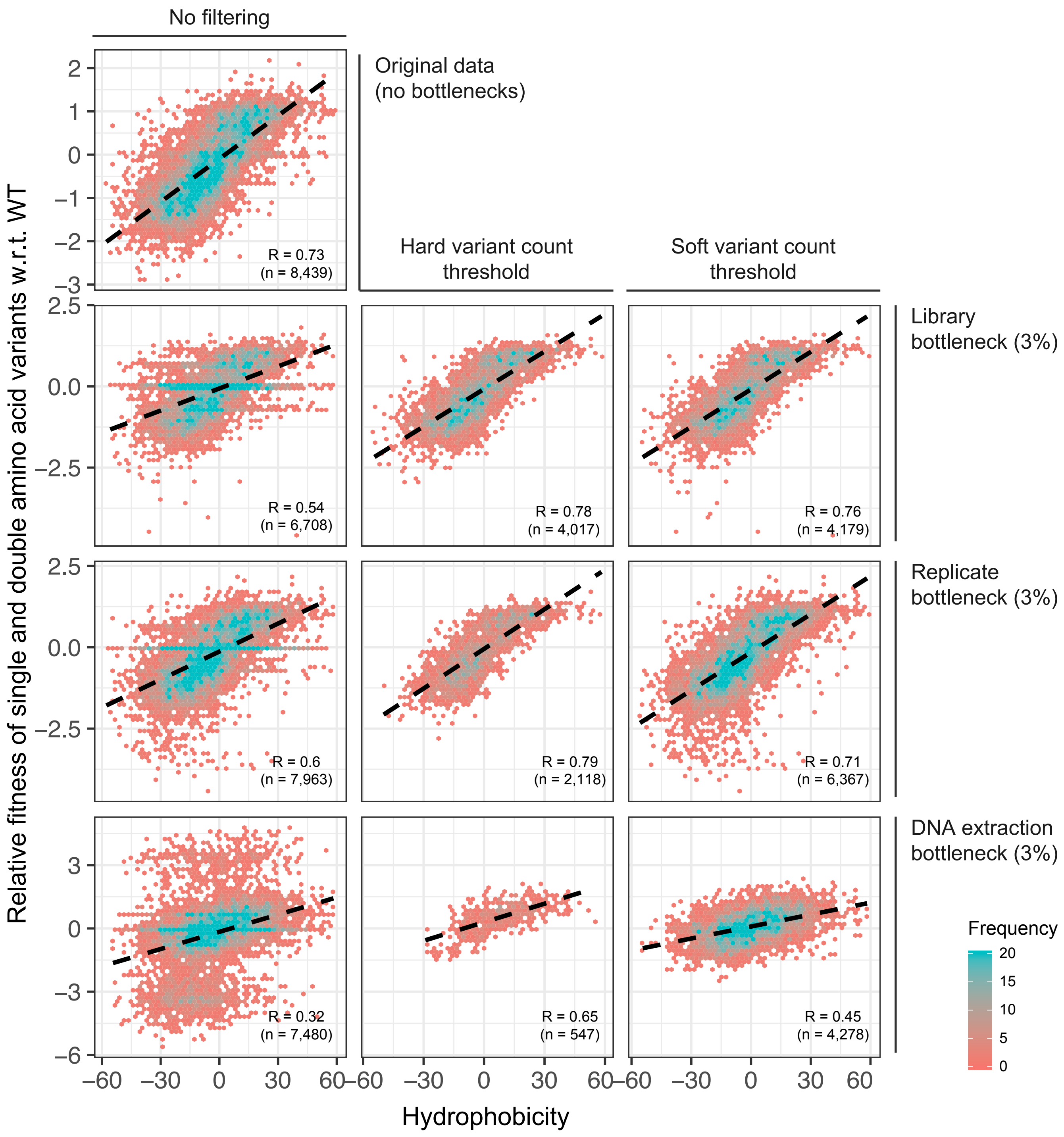


### Supplementary Figure 10
